# Material category of visual objects computed from specular image structure

**DOI:** 10.1101/2019.12.31.892083

**Authors:** Alexandra C. Schmid, Pascal Barla, Katja Doerschner

**Affiliations:** Department of Psychology, Justus Liebig University Giessen, Germany; INRIA, University of Bordeaux, France; Bilkent University, Turkey

## Abstract

Recognising materials and their properties from visual information is vital for successful interactions with our environment, from avoiding slippery floors to handling fragile objects. Yet there is no simple mapping of retinal image intensities to the physical properties that define materials. While studies have investigated how material properties like surface gloss are perceived from regularities in image structure, such as the size, sharpness, contrast, and position of bright patches caused by specular reflections, little is known how this translates to the recognition of different material classes like plastic, pearl, satin, or steel, and the underlying mechanisms involved. We investigated this by collecting human psychophysical judgments about complex glossy objects rendered in natural illumination fields. We found that variations in specular image structure – produced either by different reflectance properties or direct manipulation of image features – caused categorical shifts in material appearance, suggesting that specular reflections provide diagnostic information about a wide range of material classes, including many that should be defined by more complex scattering functions. Moreover, differences in material category were predicted by, but also appeared to mediate, cues for surface gloss, providing evidence against a traditional feedforward view of neural processing that assumes combinations of mid-level properties mediate our holistic, categorical impressions. Instead, our results suggest that the image structure that triggers our perception of surface gloss plays a direct role in visual categorisation and, importantly, that the perception and neural processing of stimulus properties should not be studied in isolation but rather in the context of recognition.

## INTRODUCTION

Our visual world is made up of light that has been reflected, transmitted, or scattered by surfaces. From this light we can tell whether a surface is light or dark, shiny or dull, translucent or opaque, made from platinum, plastic, or pearl (Figure 1A). Yet material perception is not trivial as the structure, spectral content, and amount of light reaching our eyes depends not only on surface reflectance, transmittance, and scattering properties, but also on complex interactions with the 3D shape, position, and orientation of surfaces with respect to each other, light sources, and the observer. Thus, our ability to recognise materials and their intrinsic properties serves as a compelling example of an important but unresolved challenge in visual neuroscience: How does the brain disentangle the conflated contributing factors to the retinal image to perceive our visual world?

**Figure 1.**
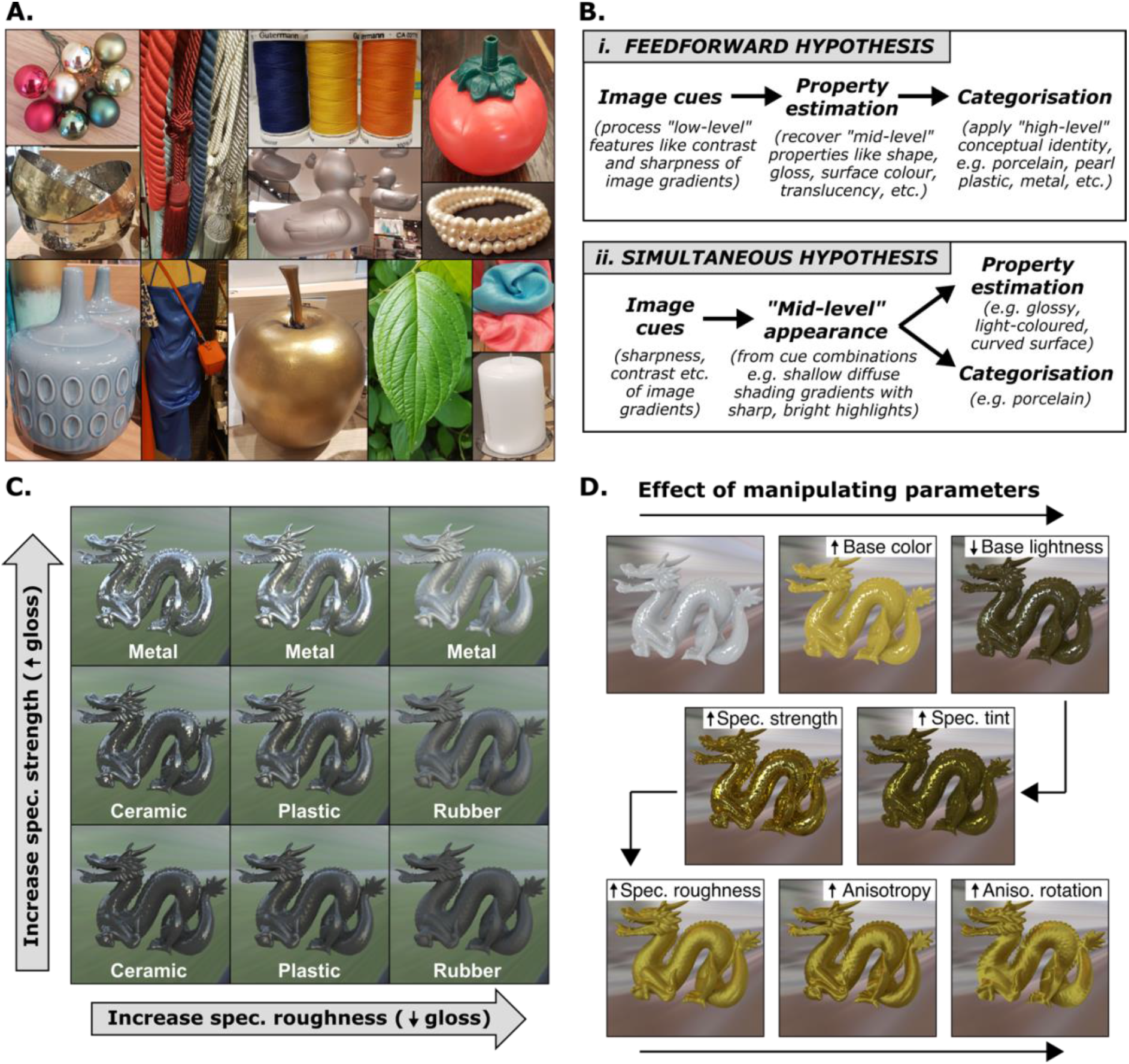
The relationship between reflectance properties, image structure, and material appearance. **A.** Most objects that we see every day are made from materials whose specular reflectance properties produce characteristic image structure. The appearance of these reflections is determined (and constrained) by the way in which these materials scatter light, in addition to other generative processes (see Figure 2). **B.** However, the extent and mechanism by which specular structure contributes to categorisation remains unknown. A feedforward view of neural processing **(i)** assumes that categories are defined by combinations of estimated mid-level properties like gloss, colour, and shape, etc., which the visual system tries to “recover” from the image (Barrow & Tenenbaum, 1978). In contrast, the simultaneous hypothesis **(ii)** assumes that the visual system naturally learns about statistical variations (regularities) in image structure, from which the identity or category of a material can be “read out” simultaneously with surface qualities like gloss (Fleming & Storrs, 2019). **C.** Some aspects of mid-level appearance (like perceived gloss) tend to correlate with physical properties (like specular reflectance), all else being equal: Systematically manipulating specular reflectance properties of otherwise identical objects causes salient visual differences in the appearance of highlights, which affects perceived gloss in predictable ways (see Figure 2). Specifically, increasing specular strength (bottom to top) increases the contrast of specular highlights, causing the dragon to appear glossier; increasing specular roughness (from left to right) decreases the clarity (or sharpness) of the specular highlights, causing the dragon to appear less glossy. These manipulations also affect our *qualitative* (i.e., categorical) impressions: the surfaces resemble different materials like glazed ceramic, glossy plastic, dull plastic, rubber, polished metal and brushed metal. Since shape, surface colour, and illumination conditions are held constant, all visual differences are caused by differences in specular reflectance properties, suggesting that specular structure may directly contain diagnostic information about material class (i.e., simultaneously with surface gloss). **D.** Reflectance parameters that were manipulated in the experiments, and examples of how this affected the visual appearance of surfaces.

While there is a growing body of work investigating the visual perception of material properties like colour, lightness, transparency, translucency, and gloss (for reviews see Anderson, 2011, 2020; Foster, 2011; Chadwick & Kentridge, 2015; Fleming 2014, 2017), there is comparatively little work investigating the recognition of different material classes like plastic, pearl, satin, steel, etc. (Balas, 2017; Baumgartner et al., 2013; Fleming et al., 2013; Lagunas et al., 2021; Nagai et al., 2015; 2018; Norman et al., 2020; Sharan et al., 2014; Tamura et al., 2018; Todd & Norman, 2018, 2019; Wiebel et al., 2013). For example, previous research has discovered a limited set of image conditions (photogeometric constraints) that trigger the perception of a glossy versus matte surface, involving the intensity, shape, position, and orientation of specular highlights (bright reflections; Beck & Prazdny, 1981; Blake & Bülthoff, 1990; Wendt et al., 2008; Todd et al., 2004; Anderson & Kim, 2009; Kim et al., 2011; Marlow et al., 2011), and lowlights (dark reflections; Kim et al., 2012) with respect to diffuse shading. However, it remains unknown what image information triggers our perception of different materials. A likely reason for this discrepancy is that studying properties like colour and gloss seems more tractable than discovering the necessary and sufficient conditions for recognising the many material classes in our environment. The challenge is that the perceptual space of materials is unspecified; there are many different optical “appearances” that can look like steel (think polished, scratched, rusted), or plastic (smooth and glossy or rough and dull).

Furthermore, a traditional feedforward view of neural processing is often assumed in which the recognition of objects and materials proceeds from the processing of *low-level* sensory information (image cues) to the estimation of shape and surface properties (often referred to as *mid-level* vision; Anderson, 2011, 2020) to the *high-level* recognition of object and material categories (*feedforward hypothesis*; Figure 1B (i); e.g., Komatsu & Goda, 2018). Within this framework it would make sense to first study how the visual system computes mid-level material properties like surface gloss and colour from images, as material class is thought to be subsequently computed from the high-dimensional feature space defined by these component mid-level properties (e.g., Fleming et al., 2013; Lagunas, 2021; Nagai et al., 2015, 2018; Schwartz & Nishino, 2020; Tanaka & Horiuchi, 2015).

An alternative view suggests that the visual system does not try recover physical surface properties like diffuse reflectance (seen as surface colour) and specular reflectance (seen as gloss) per se., but rather learns about statistical variations or regularities in image structure from which both material properties like gloss and material categories like plastic can be “read out” simultaneously (*simultaneous hypothesis*, Figure 1B (i); Fleming, 2014, 2017; Fleming & Storrs, 2019; Storrs & Fleming, 2021). For example, in Figure 1C, systematic differences in the surface reflectance properties of otherwise identical objects produce variations in the appearance of specular highlights, such as their contrast, sharpness, and coverage (see Figure 2). These variations in highlight appearance not only affect the (quantitative) level of surface gloss perceived (shiny to dull), but also the *quality* – or material category – of the surface altogether (rubber, plastic, ceramic, or metal), and it is possible that the same image features, or *cues*, directly underlie both processes. As it stands, however, the relationship between image cues, surface properties like gloss, and material recognition is unclear.

**Figure 2.**
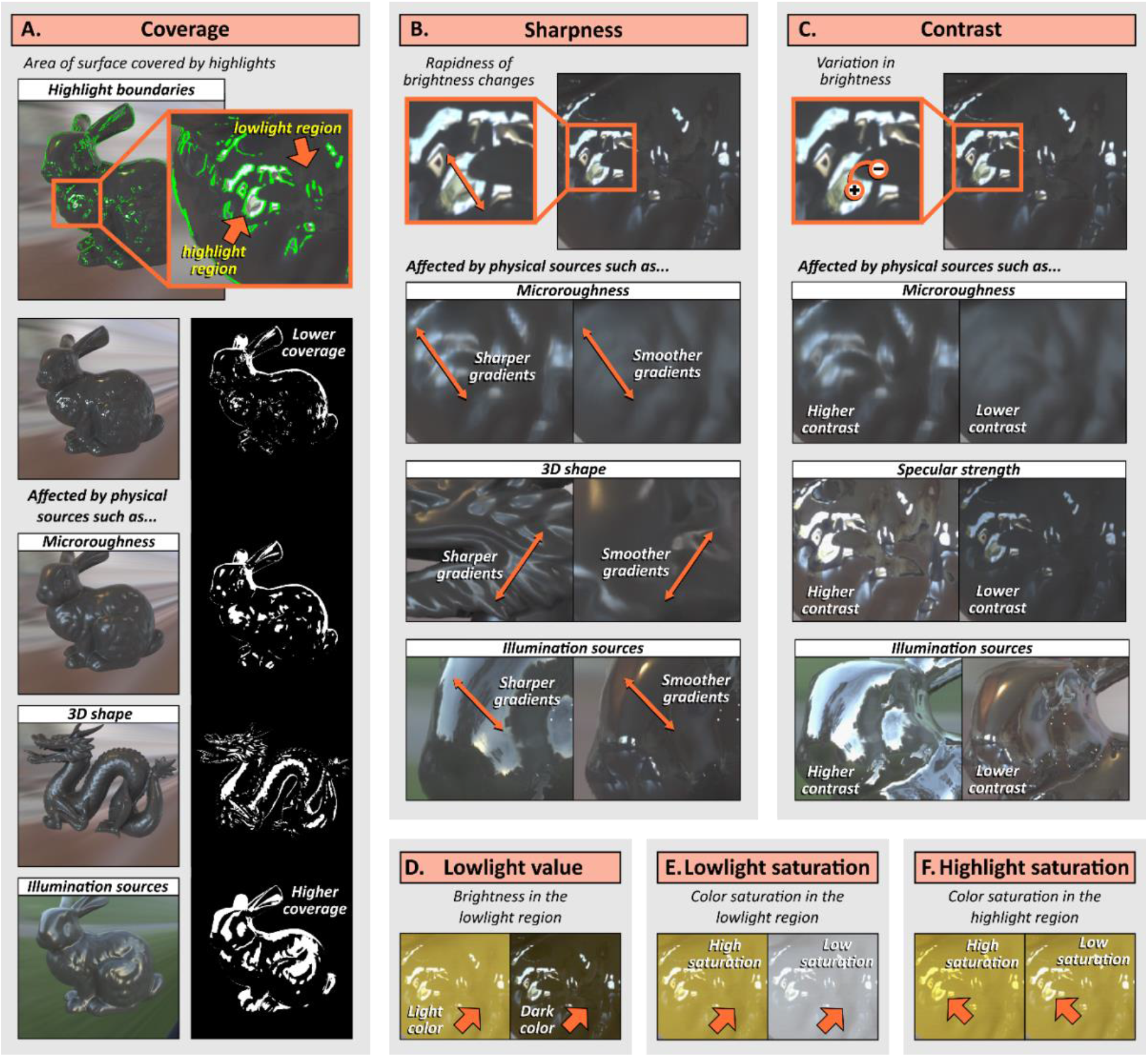
Image cues produced by reflectance properties. **A-C.** The surface appearance of glossy objects is constrained by generative processes, which determine the coverage, sharpness, and contrast of specular highlights in the image. Specifically, image structure is constrained by the way that surface reflectance properties interact with 3D surface geometry and the light field (e.g., Marlow & Anderson, 2013). *Microroughness:* Whereas matte surfaces scatter light in all directions, glossy surfaces reflect light directionally, preserving the structure of the illumination field in the resulting image. Highly smooth surfaces produce narrow specular lobes (Klinker, Shafer, & Kanade, 1988) that increase the contrast and clarity (sharpness) of the reflected image relative to microscopically rougher surfaces, which produce broader specular lobes that blur this structure. Surfaces with higher microroughness also cause specular highlights to appear more spread out (higher coverage) due to the larger range of surface normal orientations reflecting bright light sources towards the observer. *Specular strength:* While dielectric materials like plastic reflect only a proportion of light specularly, metals reflect all light specularly, increasing the contrast of the reflected image. *Illumination sources:* Because of the structure-preserving properties of glossy objects, the contrast, sharpness, and coverage of specular highlights depend on the incident light that is being reflected, i.e., the intensity and structure of the illumination field. *3D shape:* Furthermore, 3D shape distorts this illumination structure. Specular reflections cling to points of high surface curvature and are elongated along (but slide rapidly across) directions of minimal surface curvature (Kim et al., 2011; Marlow et al, 2011; Fleming et al., 2004; Koenderink & van Doorn, 1980). This means that 3D shape and observer viewpoint affect the location and distortion of specular reflections, and thus proximal stimulus properties of coverage, sharpness, and contrast of the highlights. **D-F.** The brightness and colour saturation inside and outside of the highlight region is also determined by interactions between a surface’s absorption/reflectance properties, 3D shape, and the illumination field. For example, specular reflections from dielectric materials like plastic preserve the spectral content of incident light, while coloured metals tint reflected light. However, reflections from plastic can look tinted if the prevailing illumination is coloured.

Here, we test the precise role of image structure produced by specular reflections in material recognition by rendering complex but carefully controlled stimuli in natural illumination fields, measuring and manipulating aspects of specular image structure, and collecting human psychophysical judgments about surface appearance in a series of experiments. Our results show that the visual system is highly sensitive to specular structure for material recognition, even in the absence of information from other physical sources like surface texture, transmittance, and subsurface scattering, suggesting that specular reflections play a more extensive role in visual recognition than previously appreciated. Furthermore, the data reveal that, rather than materials being derived via the estimation of material properties like gloss (*feedforward hypothesis*), it is more likely that image cues for gloss perception are constrained by material class, implying that the perception and neural processing of gloss and other stimulus properties should be studied in the context of recognition rather than in isolation. Moreover, our results demonstrate that material category is directly computable from image-measurable cues generated by surface reflectance properties and that manipulating these cues transforms perceived category, suggesting that specular structure provides direct diagnostic information about an object’s material. We discuss a simultaneous account of material perception (*simultaneous hypothesis*) in which stimulus properties like gloss are co-computed with (i.e., constrained by) our qualitative holistic impressions.

## RESULTS

### Specular reflection appearance yields a diverse range of material classes

If the visual system is sensitive to specular reflections for material recognition beyond whether a surface is shiny or matte (Marlow et al., 2011), then altering the appearance of specular reflections should lead to changes in perceived material. To test this, we computer-rendered glossy objects with different surface reflectance properties under natural illumination (Figure 1D). We parametrically manipulated base colour (lightness and saturation of the diffuse component) in addition to five specular reflection parameters (specular strength, specular tint, specular roughness, anisotropy, and anisotropic rotation) to control the appearance of specular reflections with respect to diffuse shading, resulting in 270 stimuli (Supplementary Figure 1). We collected unbiased participant-generated category terms for each stimulus using a free-naming task (Experiment 1) where participants (n=15) judged what material each object was made from with no restrictions.

After processing for duplicates and similar terminology (see Analyses), over two hundred terms were used to describe the materials, more than half of which were category terms (i.e., nouns like porcelain, gold, plastic, stone, ceramic, chocolate, pearl, soap, wax, metal, bronze, rubber, fabric, velvet, etc.; Supplementary Figures 2 and 3). Although the use of each term was distributed well among participants (i.e., category labels did not come from the same few participants), semantic labels are only relevant to the extent that they capture qualitative perceptual differences in visual material appearance. For example, dark brown stimuli with medium-clarity specular reflections might be labelled as “melted chocolate” or “mud” but would be qualitatively visually equivalent to one another. Such stimuli would have a different visual quality to light yellow stimuli with low-clarity, dim reflections, which might be labelled as “wax” or “soap” (Figure 3A). Therefore, we sought to reveal the latent perceptual space of materials for our stimulus set, with the following steps.

**Figure 3.**
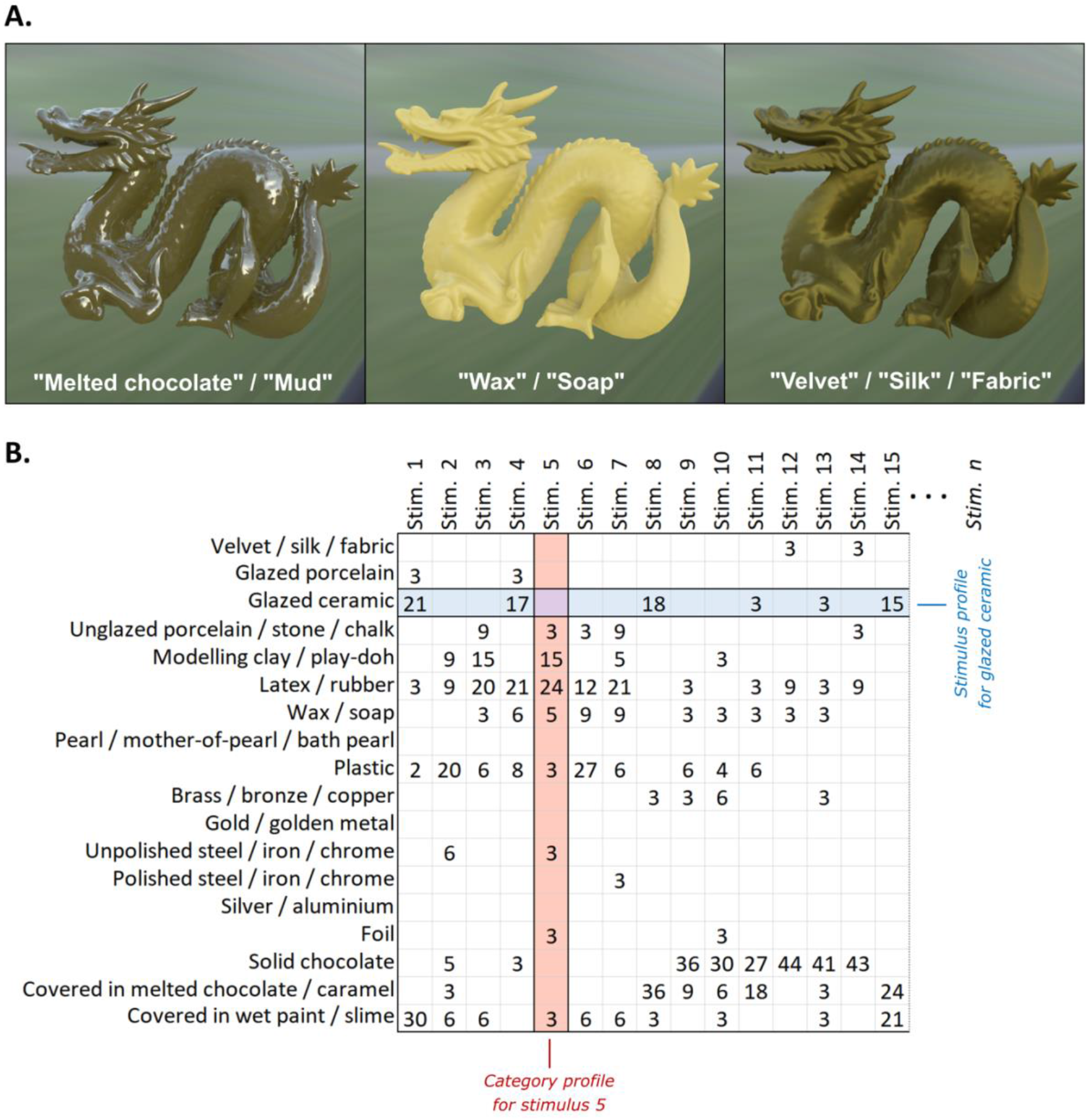
Multiple semantic terms can describe the same qualitative visual appearance of a material. **A.** The object in the first panel has a dark brown, lumpy surface with medium-clarity specular reflections and could be labelled “melted chocolate” or “mud”. This surface has a different visual quality to the object in the middle panel, which has a light-yellow surface with low-clarity, dim reflections that make it look like “wax” or “soap”. The object in the third panel has very rough, anisotropic specular reflections that rapidly change in brightness at the boundaries between highlights and lowlights, giving it the visual characteristics of “velvet”, “silk”, or “fabric”. **B.** Sum of confidence ratings from the 18-AFC experiment (Experiment 2) for each stimulus and each category for the first 15 stimuli (out of 924). *Category profiles* are the distribution of category responses for each stimulus and reveal the extent to which multiple category terms apply to the same stimulus. *Stimulus profiles* are the distribution of stimuli that were allocated to each category and reveal the extent to which category terms are correlated with one another. Stimulus profiles were used to reveal the latent perceptual space of materials for our stimuli (factor analysis; see Figure 4), and category profiles were used to calculate material dissimilarity scores between each pair of stimuli (representational similarity analysis; see Figure 7).

First, we reduced the set of category labels generated from the free-naming task to those that were used by at least five participants, and merged visually or semantically similar terms, guided by correlations between the categories (see Analyses and Supplementary Figure 4). The reduced set of 18 category terms is shown in Figure 3B.

Second, a separate set of participants (n=80) completed a multiple-alternative-forced-choice task (18-AFC task; Experiment 2) where they were asked to choose the material category that best applied to each stimulus (Supplementary Figure 5). For this experiment, the stimulus set was extended to include a larger range of reflectance parameters, two shapes (dragon and bunny), and two lighting environments (kitchen and campus), resulting in 924 stimuli (Supplementary Figure 1). Participants (n=20 per stimulus) provided confidence ratings (converted to a score between 1-3), which allowed them to indicate their satisfaction with the category options presented. The confidence ratings for each category for each stimulus were summed across all participants, providing a distribution of category responses for each stimulus (*category profiles*) and a distribution of stimuli for each category (*stimulus profiles*; Figure 3B). For many stimuli, more than one category term applied; for example, Stimulus 1 was almost equally classified as “glazed ceramic” and “covered in wet paint”, Stimulus 6 was classified as both latex/rubber and plastic, etc. (Figure 3B). This might be due to redundancies in terminology and/or imperfect category membership driving different decision boundaries (e.g., “it looks a bit like plastic but also a bit like rubber”). Indeed, we found that the stimulus profiles for some of the categories correlated with one another (Supplementary Figure 6).

Third, the data were subjected to a factor analysis to extract the common variance between categories and reveal orthogonal perceptual dimensions for our stimulus set. Figure 4A shows that there was no clear plateau in shared variance explained with each additional factor, and twelve factors were needed to account for at least 80% of the shared variance between stimuli (the upper limit based on degrees of freedom; see Analyses). Figure 4B shows example stimuli from the emergent dimensions, which were highly interpretable (twelve plus one dimension that emerged from the negative loadings; Figure 4C). Retaining eight or ten factors accounted for approximately only 60% and 70% of the common variance between categories, respectively, demonstrating that changing the appearance of specular reflections yields a diverse range of perceived materials that cannot be reduced to a small number of perceptual dimensions.

**Figure 4.**
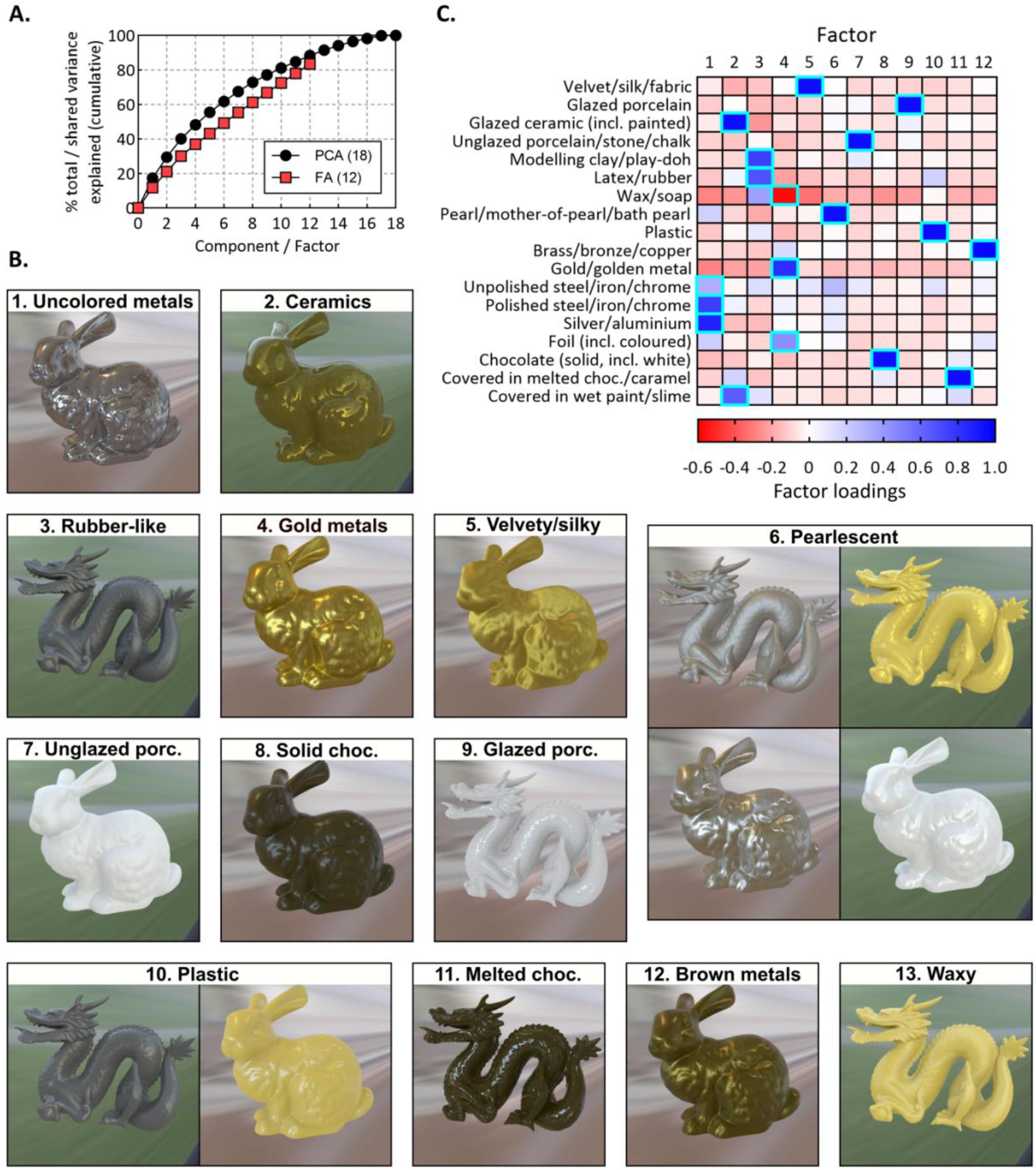
Factor analysis performed on stimulus profiles obtained from the 18-AFC task. **A.** Black circles: cumulative total variance explained by each component from an initial principal components analysis. Red squares: Cumulative shared variance explained by each factor from the 12-factor solution, which accounted for 80% of the shared variance between stimulus profiles. **B.** Example stimuli from the 13 emergent material dimensions from the factor analysis. **C.** Heat plot of the factor loadings for each category from the 18-AFC task, with blue and red cells showing positive and negative loadings, respectively. The highlighted cells show the factor onto which each category most strongly loaded. Note that the 13^th^ dimension emerged from the negative loadings onto Factor 4. The emergent dimensions were highly interpretable from the category loadings onto the factors; for example, uncoloured metals like steel and silver all loaded strongly onto factor 1, forming a single dimension. We gave material category labels to each dimension (shown in **B**); however, note that these are arbitrary and are only included for interpretative convenience. Supplementary Figure 7 shows the results of a principal components analysis that retained all dimensions in addition to the results of other factor solutions, whose dimensions overlap with those here. Supplementary Figure 8 shows more example stimuli for each shape and light field.

Surprisingly, the materials that were perceived extended beyond those expected based on the reflectance function used to generate them. In the real world, materials like porcelain, pearl, soap, wax, velvet, and many others produce image structure that is caused by the extent and way in which they transmit, internally scatter, and disperse light in addition to pigment variations and mesoscale details like fibres in fabrics, or scratches in anisotropic metals. Yet, participants reported seeing these materials for surfaces that were only defined by (uniform) diffuse and specular reflectance properties, despite the absence of these other potentially diagnostic sources of image structure. Thus, our results reveal that the human visual system is highly sensitive to the image structure produced by specular reflections for material recognition, even for complex materials.

### Material class is not determined by but may mediate gloss perception

While manipulating surface reflectance properties changed image structure in a way that participants interpreted as different materials, how this image structure relates to material recognition and the underlying mechanism is unclear. A feedforward approach assumes that the appearance of specular reflections determines surface glossiness, which in turn combines with other estimated mid-level properties (such as object shape, colour, translucency, etc.) to define the material category (Figure 1B (i)). If this is true, then we should be able to identify image cues that predict gloss perception, and the material categories from Experiment 2 should be associated with a particular level of gloss. We tested this in Experiment 3 in which a separate group of participants (n=22) rated the perceived glossiness of each of the 924 stimuli from Experiment 2. We directly measured visual features (cues) of the stimuli that describe the appearance of specular reflections and are based on generative constraints on how light interacts with surfaces that have a diffuse and glossy component (Figure 2). Three of these cues – *coverage*, *sharpness*, and *contrast* – have previously been found to predict participants’ judgments of surface glossiness (Marlow et al., 2012; Marlow & Anderson, 2013), so we refer to them as *gloss cues*. However, whereas Marlow and colleagues used perceptual judgements of each cue to predict gloss, here we operationalised the cues using objective, image-based measures (see Analyses and Supplementary Figure 9). Intuitively, *coverage* is the extent to which an object’s surface is covered in specular highlights (bright reflections; Figure 2A); *sharpness* refers to the rapidness of change between brighter and dimmer regions within and at the boundary of those highlights, and is usually related to the distinctness of the reflections (i.e., the clarity of the reflected environment; Figure 2B); and *contrast* is the variance in bright and dim regions caused by specular reflections, and usually relates to how bright the specular highlights look in relation to the surrounding regions (Figure 2C).

We found that inter-subject agreement for gloss ratings was high (median r = 0.70), and overall a linear combination of the gloss cues (coverage, sharpness, and contrast) accounted for 76% of the variance in perceived gloss (Figure 5A), R^2^ = 0.76, F(3,916) = 973.74, p < 0.001. This links perceived gloss with objective, image-computable measures of specular reflection cues for a wide range of reflectance conditions. However, the material dimensions defined in Experiment 2 were not associated with a particular level of gloss, contrary to what would be predicted by the *feedforward hypothesis* (Figure 1B (i)). Instead, stimuli from the same material class exhibited a wide distribution of gloss levels, and stimuli from very visually distinct classes like ceramic and (gold and uncoloured) metals had completely overlapping gloss distributions (histograms in Figure 5B). We investigated whether the wide gloss distributions could be caused by the continuous nature of the material dimensions (e.g., perhaps more metallic-looking materials are glossier, more rubber-looking materials are less glossy, etc.). If this is true, then perceived gloss should correlate with the loading of stimuli onto material dimensions (i.e., material factor scores). This was the case for only about half of the material dimensions (scatter plots in Figure 5B, correlation coefficients highlighted in red), with the other half showing no significant correlation between gloss level and score (black coefficients), suggesting that perceived gloss level is overall not a good indicator of how well stimuli load onto a material dimension. This raises the possibility that an object’s material is computed from specular reflection image cues directly rather than via gloss estimation, which is in line with the *simultaneous hypothesis* (Figure 1B (ii)). Indeed, material (factor) scores nearly always correlated with one or more of the image cues that predicted gloss (i.e., coverage, sharpness, and contrast; Figure 5C, left), and this relationship between image cues and material scores can explain the extent to which gloss predicted material scores in Figure 5B (shown in Figure 5D). Further supporting this idea, we found that a model predicting material score from gloss level performed worse than a model predicting material score directly from a linear combination of the image cues for most material classes (Figure 5E (i)).

**Figure 5.**
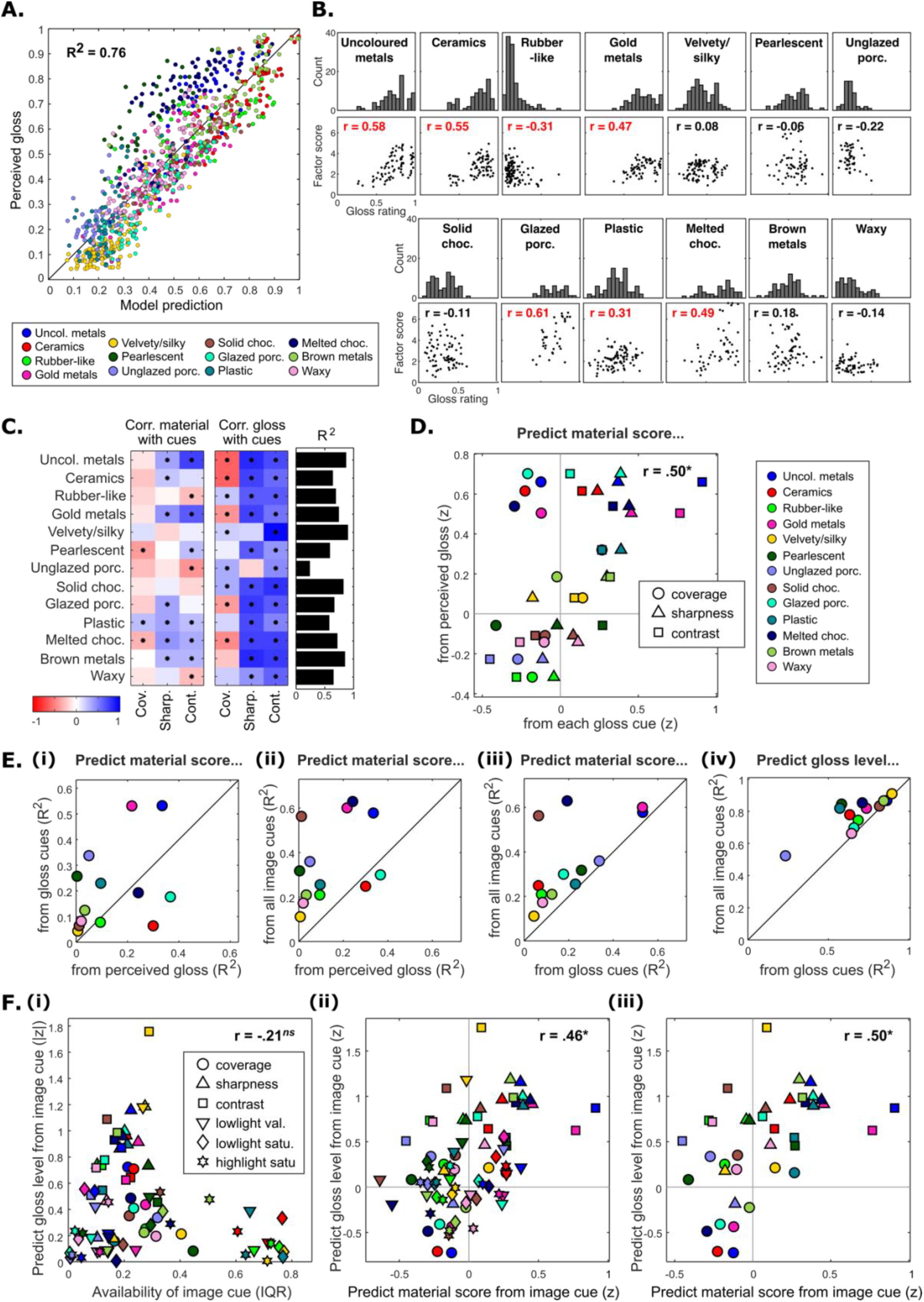
Material class is not determined by but may constrain gloss perception. **A.** Results of a linear regression model predicting perceived gloss from coverage, sharpness, and contrast (gloss cues). The model accounts for 76% of the variance in gloss ratings between stimuli and is comparable to a model that predicts participant’s gloss ratings from each other (leave one out; average R^2^ = .72). **B.** Histograms show that stimuli from the same material class exhibited a wide distribution of gloss levels, which persisted even when only the strongest loading stimuli on each dimension were considered (Supplementary Figure 10). Although some dimensions have a narrower range of gloss levels than others, stimuli from very visually distinct material dimensions like ceramics and (gold and uncoloured) metals have completely overlapping distributions of gloss ratings. Subjacent scatter plots show correlations between perceived gloss and material score, for each material class (the results hold for other factor solutions like FA8 and FA10, not shown; Supplementary Figure 11 shows (B) for PCA18 solution). **C.** The variance in gloss accounted for by the cues differed within each material class (R^2^, black bars), and the strength and direction of each cue’s correlation with perceived gloss / material score differed across materials (Pearson correlation colour coded according to strength and direction; see Supplementary Figure 12 for correlation between image cues). **D.** The correlations between perceived gloss and material score (from B) are confounded with the material scores correlating with the gloss cues themselves. **E.** Specular gloss cues predict material score (R^2^) better than perceived gloss for most categories and adding colour cues improves this (ii and iii). Furthermore, perceived gloss is sometimes better predicted with the inclusion of colour cues (iv). **F.** The predictability of an image cue for perceived gloss is unrelated to the “availability” (as measured by interquartile range; IQR) of those cues within a given class (i), but *are* related to the cues that predict material (ii – all six cues; iii – three gloss cues). In (D) and (F) z stands for Fisher-transformed correlation coefficients (Pearson correlation). Datapoints in (A), (D), (E), and (F) are colour coded by material category (legend is shown in A and D).

However, if material can be computed directly from specular image structure, this is unlikely to be limited to cues for gloss (i.e., coverage, sharpness, and contrast). Many material dimensions that emerged from our stimulus set seem to be defined by specific colour information produced by reflectance properties: for example, gold, brown, and uncoloured metals; melted and solid chocolate; glazed and unglazed porcelain; and waxy materials (Supplementary Figure 8). We measured three *colour cues* – *highlight saturation*, *lowlight saturation*, and *lowlight value* – that describe colour variations in specular image structure that arise from generative processes (i.e., how light interacts with diffuse and specular reflectance properties; Figure 2D-F). Intuitively, *highlight saturation* is the colour saturation within the specular highlight regions (bright spots); *lowlight saturation* is the colour saturation outside of the highlight regions (referred to as lowlight regions; Kim et al., 2012); and *lowlight value* is the brightness of the colour within the lowlight regions. Figure 5E (ii) and (iii) show that the addition of the colour image cues led to an overall improvement in predicting material scores.

Collectively, these results do not support the idea that gloss mediates the perception of materials from specular reflection image cues (*feedforward hypothesis*; Figure 1B (i)) but are in line with the idea that surface gloss and material class could be simultaneously “read out” from these cues (*simultaneous hypothesis*; Figure 1B (i)). One potential mechanism for this is that image cues independently determine gloss (gloss cues) and material (gloss cues and colour cues). However, Figure 5E (iv) shows that, in addition to gloss cues, colour cues also contribute to perceived gloss for some materials, suggesting that cues for gloss and material might not be independent. In fact, the contribution of image cues to perceived gloss was not stable across different material classes (black bars in Figure 5C), as would be expected if the visual system computed surface gloss from image cues independent of material class. Instead, the predictiveness of each cue to gloss differed with material (Figure 5C, middle), and this covaried with how well each cue predicted material score (that is, cues that predicted gloss also tended to predict material score; Figure 5F (ii) and (iii)). This dependency between cues for gloss and cues for material is unlikely to be due to gloss mediating cues to material (*feedforward hypothesis*) as our previous analyses showed that gloss level was not a good indicator of material class. Nor can the differing contributions of cues to gloss be explained by the relative availability of those cues within each class (i.e., cue variability; Figure 5F (i)). This implies that, rather than gloss mediating cues to material, the image cues used to estimate gloss could instead be mediated by perceived category. This does not necessarily contradict the *simultaneous hypothesis*, as it is possible for surface gloss and material to be co-computed and mutually constrained by the same image structure (i.e., computed *simultaneously* but not *independently*). We elaborate upon this idea in the General Discussion, but first test the important prediction that material categories can be discriminated directly from features of specular image structure.

### Material class is predicted directly by features of specular image structure

To test the prediction that materials can be discriminated directly from features of specular image structure (*simultaneous hypothesis*; Figure 1B (ii)), we sought to predict material class from the measured image cues (Figure 2) for the stimuli that loaded most strongly onto each material dimension (Figure 4). Figure 6A plots the distribution of image cues for these stimuli (coloured violin plots) for each material. Visualised in this way, the image cues provide “material signatures” for each class. The data were subjected to a linear discriminant analysis (LDA) that classified materials based on linear combinations of image cues. We used a leave-one-condition-out approach, where the classifier was trained on three out of the four shape/lighting conditions (e.g., dragon-kitchen, bunny-kitchen, dragon-campus) and tested on the remaining condition (e.g., bunny-campus). Figure 6B plots the accuracy of the model for each material, combined over the four training-test combinations. Overall accuracy was 65%, which is well above chance (7.7%, red dotted line). A further cross-validation test showed that the model generalised across shape and lighting conditions (Figure 6C), demonstrating that feature of specular image structure can predict human material categorisation behaviour for our stimulus set. Figure 6D and E illustrate that the discriminations made by the model are perceptually intuitive.

**Figure 6.**
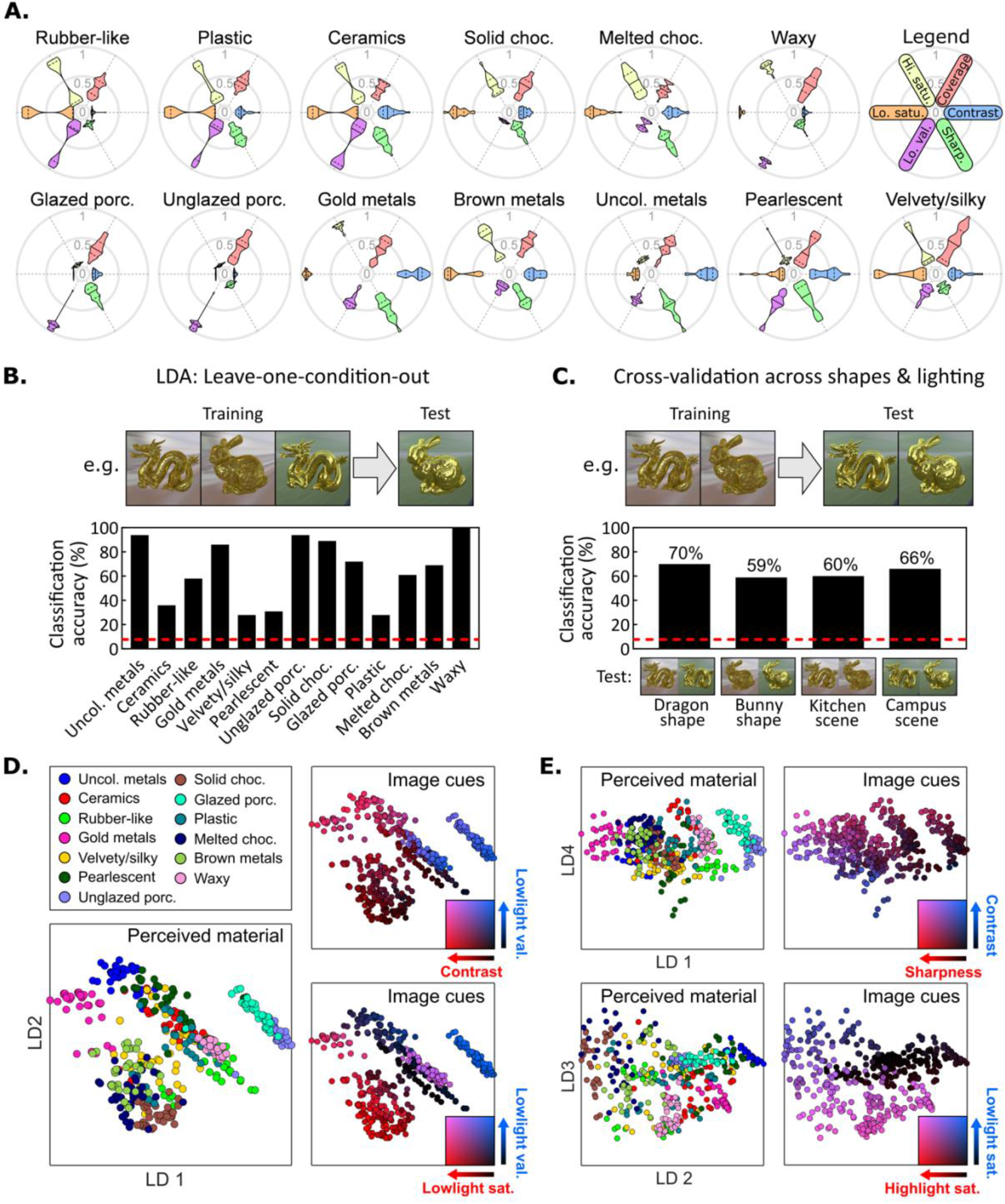
Linear discriminant analysis (LDA) predicting material category from image cues. **A.** The radial violin plots show the distribution of measured image cues for each material. For each cue, solid lines correspond to the 50^th^ percentile and dashed lines to the 25^th^ and 75^th^ percentile of the distribution. See text for details. **B.** The results of a linear classifier with leave one condition out validation procedure. The red dotted line indicates chance level (1/13). **C.** Classification accuracy generalises across different illumination and shape conditions. **D.** The stimuli are plotted in linear discriminant space (LD1 and LD2 stand for linear discriminant 1 and 2, respectively). Points are colour coded by either category (left), or image cues (right). **E.** The same stimuli are plotted for different linear discriminants (top: LD1 vs. 4; bottom: LD 2 vs. 3. These plots illustrate that the discriminations made by the model are perceptually intuitive: For example, a combination of average saturation and brightness within the lowlight region can be used to discriminate materials like porcelain (bright, uncoloured body) and waxy materials (bright, coloured body) from other materials with darker body colours; saturation within the highlight region is useful for discriminating brown metals (coloured highlights) from solid chocolate (uncoloured highlights). See Supplementary Figure 13 for similar results with other dimension reduction solutions.

Interestingly, some materials were classified better than others (Figure 6B). The materials with the highest classification accuracies (metals, chocolate, porcelain, and waxy materials) were those with the most distinct material signatures (Figure 6A). That is, there are at least a few features that seem to “characterise” those materials (e.g., silvers must have uncoloured highlights and lowlights; wax must be light and coloured with low-contrast reflections). The materials with the lowest classification accuracies (ceramics, pearlescent materials, plastic, and fabrics) had a less distinct set of image cues. One possible reason for this is that there are, for example, many types of plastics or pearlescent materials (Figure 4B), and different specific (nonlinear) combinations of cues define these subtypes – something that LDA does not capture. A second possibility is that the stimuli might not fall nicely into perceptually discrete classes and would be better represented as a smooth continuation from one material dimension to another. To allow for this, we next tested whether variations in category profile between stimuli (Figure 3B) could be predicted by variations in the measured image cues (Figure 2) using representational similarity analysis (RSA; Kriegeskorte et al., 2008). RSA additionally allows for direct comparison between image cues for material discrimination and gloss perception.

### Material differences are better predicted by image cues for gloss than by gloss itself

If the same features of specular image structure underlie material discrimination and gloss perception (*simultaneous hypothesis*; Figure 1B (ii)), then perceived differences in both material (Experiment 2) and gloss (Experiment 3) should be reflected by differences in the measured colour and gloss cues (Figure 2). Furthermore, differences in material should not be better predicted by perceived gloss (*feedforward hypothesis*) than by image cues for gloss directly. To test these predictions, we calculated dissimilarity scores for each pair of stimuli in terms of perceived material, perceived gloss, and measured image cues (Figure 7A). Material dissimilarity scores were calculated as one minus the spearman correlation coefficient for each pair of category profiles. Gloss dissimilarity scores were calculated as the absolute difference between average gloss ratings between each stimulus pair, and dissimilarity scores for each image cue were calculated in the same way (using absolute difference). Each point in the matrices (known as representational dissimilarity matrices or RDMs; Kriegeskorte et al., 2008) shows how dissimilar each stimulus is to each other stimulus in terms of their category profile, gloss ratings, and image cues, with yellow pairs being more dissimilar.

Dissimilarity scores in the upper triangles of each RDM (highlighted in red) were vectorized and subjected to two linear regressions that predicted material dissimilarity and gloss dissimilarity (separately) from image cue dissimilarities (Figure 7B). Figure 7C shows that image cue dissimilarities significantly predicted differences in both perceived material, R^2^ = 0.32, F(6,426419) = 32797, p < 0.001, and perceived gloss, R^2^ = 0.42, F(6,426419) = 50635, p < 0.001. These image-cue models performed slightly better than models using differences in reflectance parameters as predictors (Figure 7C), demonstrating that the image cues capture the variance explainable (by a linear model) in material and gloss caused by changes in surface reflectance. The slight improvement in predictive value for image cues over reflectance parameters is likely due to the fact that the cues additionally account for differences in image structure caused by shape and lighting differences, which can affect material appearance (e.g., Norman et al., 2020). Furthermore, we found that the model containing (all six) image cues (R^2^ = 0.32) predicted changes in material better than a model containing only colour cues (lowlight value, lowlight saturation, highlight saturation) in conjunction with perceived gloss, R^2^ = 0.23 (Figure 7C), providing further evidence that material recognition occurs directly from image structure that (simultaneously) triggers our perception of mid-level properties like gloss, rather than via estimation of those properties in a feedforward manner.

**Figure 7.**
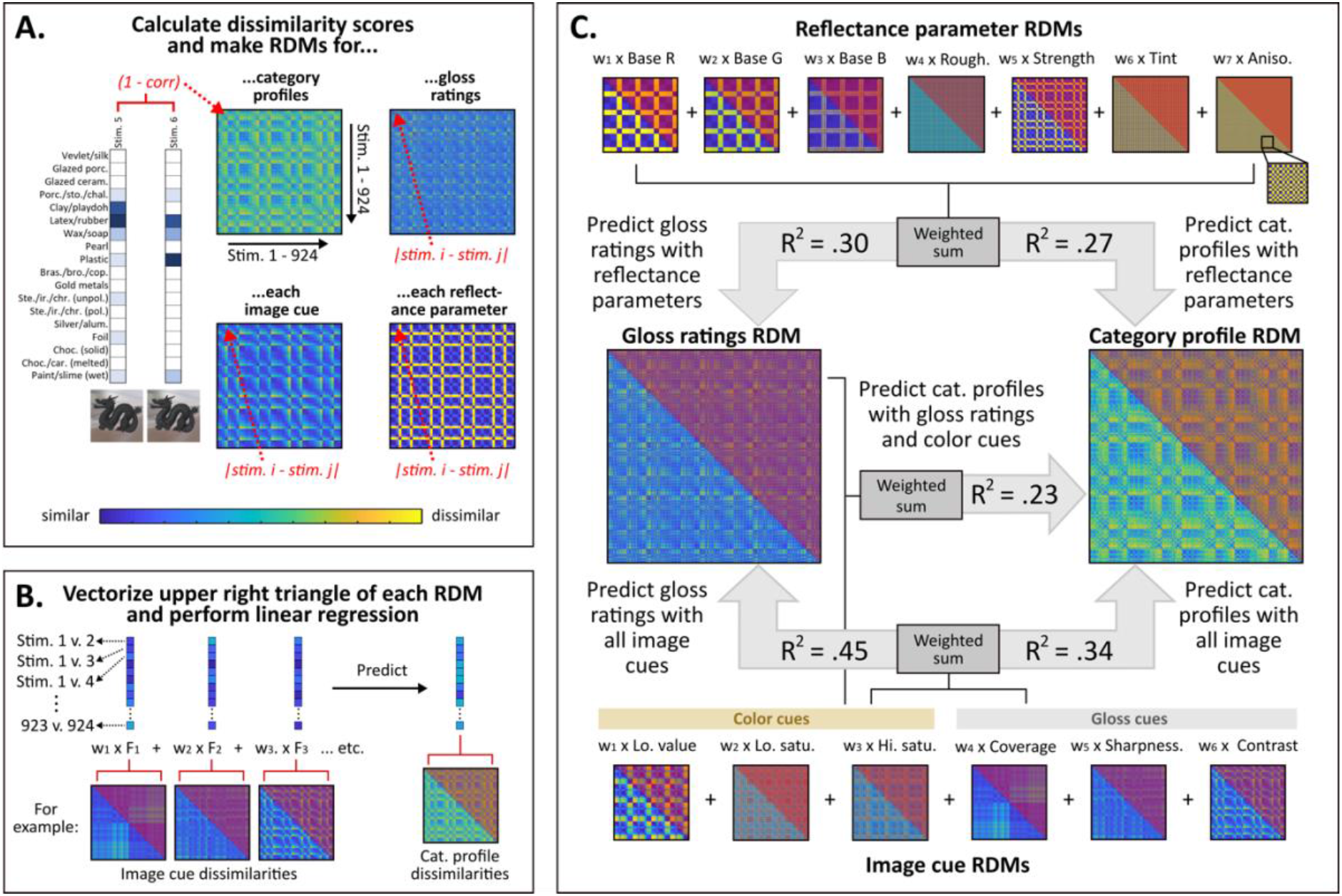
Representational similarity analysis (RSA) predicting dissimilarities in material and gloss from image cue differences. **A.** Representational dissimilarity matrix (RDM) for category profiles. Each point in the matrix shows the dissimilarity between two stimuli, calculated as one minus the Spearman correlation (*r*) of their category profiles. We also computed RDMs for average gloss ratings, each of the six image cues, and each reflectance parameter (7), **B.** Dissimilarity scores in the upper triangles of each RDM (highlighted in red) were vectorized and subjected to linear regressions that predicted material dissimilarity and gloss dissimilarity (separately) from image cue dissimilarities (panel C, bottom) and reflectance parameter dissimilarities (panel C, top). All RDMs were normalised between zero (most similar) and one (most dissimilar). **C.** The results of each regression analysis. Models based on image cue dissimilarity outperformed those based on reflectance parameter dissimilarity for both gloss and material.

### Direct manipulation of specular image structure transforms material class

Thus far our analyses have been correlational in nature. If material class is computed from combinations of the image cues that we measured (Figure 2), then directly manipulating these cues should transform perceived category in predictable ways. This would better test whether the measured image cues are causally responsible for the perceived category shifts, or whether they merely correlate with other changes in image structure that are important for material perception that we did not measure.

To this end, we attempted to directly manipulate image cues to transform stimuli that were perceived as glazed ceramic in Experiment 2 into each of the remaining materials, for each shape and light field (Figure 8A). We created two stimulus sets to test the success of our cue manipulations. The first set contained “simple” (linear) image cue manipulations, which closely correspond to the previously measured image cues (see Supplementary Figure 14). The second set contained “complex” (nonlinear) image cue manipulations, which accounted for particular types of contrast needed for some materials that we empirically noticed were not captured by the first set of manipulations. For example, the velvety/silky stimuli from Experiment 2 were exclusively defined by very rough, anisotropic specular reflections, which caused a high degree of directional blur. This gives the effect of elongated, low-clarity specular reflections that, despite this low clarity, have rapidly changing (sharp/high contrast) boundaries between highlights and lowlights relative to isotropic surfaces (see Figure 3A, Figure 4B, and Supplementary Figure 8). On the other hand, the perception of uncoloured metals appears to require the presence of high contrast reflections that appear all over the surface (i.e., there is high contrast image structure in the lowlights, not just the highlights; see also Norman et al., 2020). These particular types of contrast could not be achieved through a simple linear transformation of image intensities from the glazed ceramic stimuli but were accounted for after applying more complex additional filters to non-linearly manipulate pixel intensities (Figure 8B and Supplementary Figure 15).

**Figure 8.**
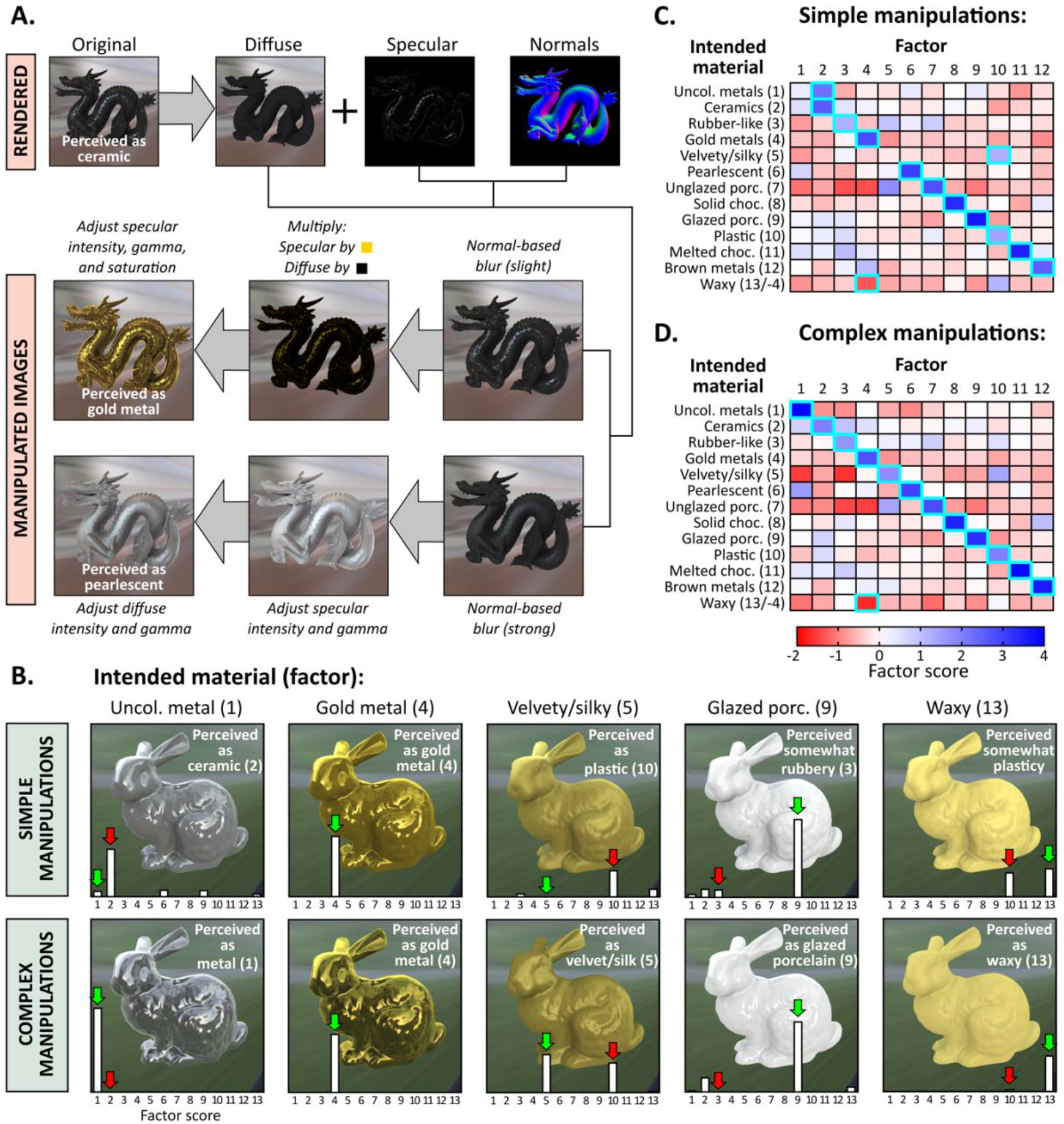
Manipulated image cues transforms perceived material category. **A.** The manipulations were performed on specular and diffuse images separately before being recombined: as with the image cue measurements, the specular component was obtained by subtraction of the diffuse component from the full rendering (see Supplementary Figure 9). The sharpness of reflections was first modified by blurring pixels of the specular component that have sufficiently similar normals (see Methods). The subsequent filters adjust the colour and intensity of each component. For instance, for materials like gold (top manipulated image), the specular component is first multiplied by a colour (the diffuse component is multiplied by 0); then its intensity and saturation are adjusted. For materials like pearl (bottom manipulated image), the intensity of the diffuse component is also adjusted. See the Methods section for a detailed description of each filter, along with the special filter used for velvet, based on a non-monotonic remapping of the specular component. **B.** Example stimuli after simple manipulations (top) and complex manipulations (bottom). White bars show the stimulus factor scores after participant judgments (see main text). After applying simple image manipulations, materials sometimes resembled unintended categories (red arrows). The perceived materials align much better with the intended category (green arrows) after the complex image manipulations. **C.** and **D.** Heat maps showing average factor scores for stimuli from each intended category after simple and complex cue manipulations, respectively. The highlighted cells show the factor onto which each category most strongly loaded.

In Experiment 4, a new set of participants (n=22) performed an 18-AFC task identical to Experiment 2, but with the image-transformed stimuli. The stimulus profiles were converted to factor scores to see which material dimensions best applied to the new stimuli. The average factor scores plotted in Figure 8C and D show that participants agreed with our informal observations. For the first stimulus set (linear image manipulations; Figure 8C), stimuli that were intended to be silver metals and fabric actually had the quality of glazed ceramic and plastic, respectively. For the second stimulus set (non-linear image manipulations; Figure 8D), the perceived materials align much better with the intended classes.

## DISCUSSION

In the present study we showed that the image structure produced by specular reflections not only affects how glossy a surface looks; it can also determine the quality, or category, of the material that is perceived. Critically, specular reflections provide more information than just distinguishing between surfaces that are matte versus glossy (Marlow et al., 2011), or plastic versus metal (Todd & Norman, 2018); we found that the perceptual dimensionality of glossy surfaces defined by a diffuse and specular component is much larger than has previously been suggested (e.g., Pellacini et al., 2000; Vangorp et al., 2017; Toscani et al., 2020; Hunter, 1937). This is likely because prior studies on gloss perception have manipulated stimulus parameters within a narrow range and used restricted tasks; for example, stimuli are simple, smooth shapes and/or their reflectance properties only encompass the range of plastic- and ceramic-looking materials, and participants judge the level of gloss or relative similarity between surfaces. Here, we used complex shapes, manipulated reflectance parameters within a wider range, and asked participants to judge each object’s qualitative material appearance. This greatly expanded the number of dimensions required to account for perceptual differences between glossy surfaces relative to prior studies. Note that we did not sample the whole perceptual space of glossy objects defined by a diffuse and specular component (e.g., we manipulated colour saturation within only one hue). The dimensionality of gloss appearance is likely to expand further upon sampling a wider range of reflectance parameters, shapes, mesostructure detail, and environment lighting conditions.

Importantly, we found that changes in specular structure – caused by either generative sources or direct image manipulation – led to qualitative shifts in material appearance beyond those expected by the reflectance function used. This demonstrates that features of specular image structure can be diagnostic for recognising a wide range of materials. A potential reason for this is that a material’s surface reflectance properties create some of its most salient optical characteristics, and, since most surfaces reflect some light specularly, relying on such characteristics could carry ecological importance when other cues to material are not available. For instance, the visual effects of translucency can be greatly diminished with frontal lighting (Xiao et al., 2014); in such cases, image cues caused by specular reflections might remain diagnostic of translucent materials (e.g., porcelain). Similarly, meso-scale details like fibres on cloth or scratches on metal might not be visually resolvable when seen at a distance; yet such materials might still be recognised from specular image structure. Indeed, these additional sources of image structure (from mesostructure or translucency) are absent from the stimuli used in the present study, and although such details might render the materials more compelling (when say compared to a photograph), the stimuli nevertheless convincingly resemble silk-like, porcelain-like, wax-like, brushed metal-like, etc. materials. Interestingly, these results are in line with findings from the computer graphics literature that show that visual effects from different types of fabrics come predominantly from specular reflections, and diffuse reflection and shadowing-masking play a much less pronounced role even for relatively matte-looking fabrics (Irawan & Marschner, 2012).

Our data do not support the notion of a feedforward path to recognition whereby the visual system combines estimates of physical properties that are first “recovered” from images; instead, our results are in line with the idea that vision assigns perceptual qualities to statistically varying image structure (Anderson, 2011; Fleming, 2012, 2014, 2017; Purves, Morgenstern, & Wojtach, 2015). Specifically, we found that gloss was not a good indicator of a material’s class; instead, materials were differentiated directly based on image-measurable specular reflection cues, suggesting that material qualities like gloss are a perceptual outcome with – rather than a component dimension of – our holistic impressions. Indeed, the perception of material qualities like gloss might even be influenced by these holistic impressions, as suggested by the fact that the contribution of different cues to gloss was not stable across different materials, but co-varied with the cues to material class. That is, the regions of feature space occupied by different material classes (i.e., different qualitative appearances) seemed to mediate the processing of those same features when estimating surface glossiness.

One potential mechanism is that emergent categories from our stimulus set could reflect cognitive decisions, and this cognitive interpretation has a “top-down” influence on which cues are used to judge surface gloss. For example, “chocolate” and “plastic” could have similar contrast and sharpness of specular highlights (similar “gloss types”) and the different labels might result from a cognitive decision by participants based on body colour (brown versus yellow). However, we cannot think of a principled reason why a cue’s influence on material category would affect how people choose to use that cue for gloss judgments.

Furthermore, we argue that such a clear perceptual scission of image structure into different layers (e.g., a gloss layer and a body colour layer) and then subsequent “cognitive reassembly” is unlikely, especially for the complex-shaped stimuli and wide sampling of reflectance parameters used in the present study. This kind of layered appearance of gloss, which can be construed as a form of transparency perception, likely only applies to smooth surfaces with little variation in surface curvature, like spheres (Kim et al., 2012). For the present stimuli, specular image structure seems to interactively combine with diffuse shading to create a gestalt-like quality for each material, such that we may not even have perceptual access to individual cues (e.g., see Supplementary Figure 1). Like with object recognition, this material quality can be given a label like “gold” or “pearl”, but nonetheless reflects a holistic impression, rather than cognitive combination of cues (also see Okazawa, Koida, & Komatsu, 2011). Such holistic impressions are likely why we could successfully manipulate image cues to transform one category into another (Experiment 4), but why linear models like LDA, and RSA (Experiment 2) did not better account for the data.

We propose an alternative mechanism to explain the co-variation between cues for surface gloss and material class, inspired by converging evidence from independent lines of psychophysical and neuropsychological research that suggests computations of stimulus properties are inherently *coupled* (Anderson, 2020; Pasupathy et al., 2019). Specifically, monkey physiology studies have found that neurons in primate area V4 respond only to specific combinations of texture and shape but not to these properties separately, suggesting joint representations of shape and surface properties (for a review see Pasupathy et al., 2019). In line with this, a series of human psychophysics studies have demonstrated that percepts of 3D shape and surface properties (e.g., gloss or translucency) are mutually constrained by specific image gradient-contour relationships (i.e., the cues for each are not separate; for a review see Anderson, 2020). Similarly, transparency impressions triggered by low contrast centre-surround displays are mutually constrained with the perception of surface lightness: a medium-grey patch surrounded by a slightly darker (homogenous) surface can appear whiteish with the quality of a flimsy transparent sheath, which occurs to the extent that luminance within the central patch is attributed to a foreground (patch) or background (surround) layer (Schmid & Anderson, 2017). In the present study, the co-variation between cues for gloss and material class could also reflect a mutual dependency (i.e., their computations could be coupled). We suggest that the perceptual attribution of specular image structure to a surface’s qualitative material appearance constrains which image features are perceived as specular reflections (versus say bright pigment or illumination, or 3D shape; Mooney & Anderson, 2014; Wijntjes et al., 2012), and thus which cues are *perceptually available* to make a surface look “glossy” or “shiny”.

Interactions between categorical material impressions and the perception of different stimulus properties warrants further investigation because the underlying mechanism will influence how we study perception in multiple domains. Within material perception, if seemingly distinct perceptual dimensions are actually intimately linked and constrained by the same image structure, this would suggest that the perception and neural processing of properties like surface gloss should in fact be considered in the context of material recognition. For example, neurons in the lower bank of the superior temporal sulcus (STS) in monkeys have been shown to preferentially respond to specific combinations of manipulated gloss parameters (Nishio et al., 2012). It is possible that different categorical material impressions are triggered by these specific combinations, driving this response.

Moreover, research into object recognition (Grill-Spector & Weiner, 2014) often investigates how visual features like texture (Long, Yu, & Konkle, 2018) and form (Bracci, Ritchie, & Op de Beeck, 2017) influence object category representations, but thus far have neglected the role of surface properties in recognition, for example by not controlling for changes in surface properties (Kaiser, Azzalini, & Peelen, 2016; Bracci et al., 2017, Zeman et al., 2019), or “scrambling” the images in a way that breaks the perception of surfaces (Long, Yu, & Konkle, 2018). Our results suggest that the specific photogeometric constraints on image structure that trigger our perception of surface gloss play an important role in visual categorisation, and that a fruitful direction for neuropsychological research could be to focus on identifying the neural mechanisms that represent objects holistically (Schmid & Doerschner, 2019).

However, we think that this role extends beyond object identification. Just as surface gloss is not a component dimension of material perception, material perception is not merely a component dimension of scene perception. A material’s qualitative appearance is useful for different aspects of navigation, such as identifying where we are (beaches have sand and water), choosing which paths to take (icy vs. muddy vs. concrete), and deciding which obstacles to avoid (solid rock vs. flexible vegetation). We also need material perception for locating items among clutter (finding a metallic pot in the cupboard); evaluating an object’s function, usefulness, or motor affordances (Can I eat it? Is it expensive or fake? How do I pick it up?), and predicting tactile properties and future states of objects (Is it heavy, sticky, or wet? Will shatter, stretch, or bounce?). Our results shed light on how image structure is transformed into our representations of surfaces with particular material “appearances”, thereby making an important contribution towards bridging the gap between “lower-level” information processing and behavioural goals such as categorisation, action, and prediction (Malcolm et al., 2016; Groen et al., 2017).

## METHODS

### Stimuli

#### Stimulus generation: free-naming, 18-AFC, and gloss rating experiments (Experiments 1, 2, and 3)

We generated our stimulus set by computer rendering complex glossy objects under natural illumination fields. Object meshes were the Stanford Bunny and Dragon from the Stanford 3D Scanning Repository (Stanford University Computer Graphics Laboratory; http://graphics.stanford.edu/data/3Dscanrep/). Wavefront .obj files of these models were imported into the open-source modelling software Blender (v2.79) and rendered using the Cycles render engine, which is an unbiased, physically based path-tracing engine.

Stimuli were illuminated by the “kitchen” and “campus” light fields from the Debevec Light Probe Image Gallery (Debevec, 1998), and the resulting scenes were rendered at a resolution of 1080 × 1080 pixels. Interactions between light and surfaces were modelled using the Principled BSDF shader, which is based on Disney’s principled model known as the “PBR” shader (Burley, 2012). The Principled BSDF shader approximates physical interactions between illuminants and surfaces with a diffuse and specular component (for dielectric materials), and a microroughness parameter that controls the amount of specular scatter. An advantage of the Principled BSDF shader over other models like the Ward model is that it accounts for the Fresnel effect. The microfacet distribution used is Multiple-scattering GGX, which takes multiple bounce (scattering) events between microfacets into account, giving energy conserving results. Although there are many parameters to adjust in the Principled BSDF shader, we manipulated only the following (details can be found at https://docs.blender.org/manual/en/dev/render/shader_nodes/shader/principled.html):

- *Base Color:* Proportion of light reflected by the R, G, and B diffuse components.
- *Specular:* Amount of light reflected by the specular component. The normalised range of this parameter is remapped linearly to the incident specular range 0-8%, which encompasses most common dielectric materials. Values above 1 (i.e., above 8% specular reflectance) are also possible. A value of 5 (the maximum value used) translates to 40% specular reflectance.
- *Specular tint:* Tints the facing specular reflection using the base colour, while glancing reflection remains white.
- *Roughness:* Specifies microfacet roughness of the surface, controlling the amount of specular scatter.
- *Anisotropic:* Amount of anisotropy for specular reflections. Higher values give elongated highlights along the tangent direction. This direction was set to radial for our stimuli.
- *Anisotropic rotation:* Rotates the direction of anisotropy, with a value of 1.0 indicating a rotation of 360°.

#### Rendering parameters: free-naming experiment (Experiment 1)

A total of 270 stimuli were generated for the free-naming experiment. Stimuli were generated by rendering combinations of the parameters for a dragon model embedded in the kitchen light field:

- *Base Color:* Six diffuse base colours based on two levels of saturation (greyscale, saturated yellow) and three levels of value (dark, medium, and light). For the greyscale stimuli, RGB values were equal for each lightness level (RGB = 0.01, 0.1, and 0.3 for dark, medium, and light stimuli, respectively). The RGB values for saturated stimuli were [0.01, 0.007, 0.001] for dark stimuli, [0.1, 0.074, 0.01] for medium stimuli, and [0.3, 0.221, 0.03] for light stimuli.
- *Specular:* Three specular levels: 0.1, 0.3, and 1, which correspond to 0.8%, 2.4%, and 8% specular reflectance, respectively.
- *Specular tint:* Two specular tint levels for saturated stimuli: 0 (no specular tint) and 1 (full specular tint).
- *Roughness:* Four roughness levels: 0, 0.1, 0.3, and 0.6.
- *Anisotropic:* Two anisotropic levels for stimuli with roughness levels greater than zero: 0 and 0.9.
- *Anisotropic rotation:* Two anisotropic rotations for anisotropic stimuli: 0 (no rotation) and 0.25 (90° rotation).

#### Rendering parameters: 18-AFC and gloss rating experiments (Experiments 2 and 3)

A total of 924 stimuli were generated for the 18-AFC and gloss rating experiments. Stimuli were generated by rendering combinations of the parameters for two shapes (dragon, bunny) and two light fields (kitchen, campus):

- *Base Color:* Six diffuse base colours based on two levels of saturation (greyscale, saturated yellow) and three levels of value (dark, medium, and light). For the greyscale stimuli, RGB values were equal for each lightness level (RGB = 0.01, 0.03, and 0.3 for dark, medium, and light stimuli, respectively). The RGB values for saturated stimuli were [0.01, 0.007, 0.001] for dark stimuli, [0.03, 0.022, 0.003] for medium stimuli, and [0.3, 0.221, 0.03] for light stimuli.
- *Specular:* Four specular levels for the darkest four diffuse shading levels: 0.1, 0.3, 1, and 5, which corresponds to 0.8%, 2.4%, 8%, and 40% specular reflectance, respectively. The light-coloured stimuli were only rendered with the first three specular levels.
- *Specular tint:* Two specular tint levels for saturated stimuli: 0 (no specular tint) and 1 (full specular tint).
- *Roughness:* Three roughness levels: 0, 0.3, and 0.6.
- *Anisotropic:* Two anisotropic levels for stimuli with roughness levels greater than zero: 0 and 0.9.
- *Anisotropic rotation:* Two anisotropic rotations for anisotropic stimuli: 0 and 0.25.

#### Stimulus generation: cue manipulation experiment (Experiment 4)

We generated the feature-manipulated stimulus set with an approach similar to compositing techniques used in visual effects. For our manipulation we need several image components of an input scene: a full rendered image, an image of the object rendered with diffuse shading only (diffuse component), a binary mask image, and an image of the surface normals. All of these components were rendered with the Cycles engine in Blender: A manipulated image is obtained via a sequence of steps that are depicted in Figure 8A: a specular component is first obtained by subtracting the diffuse component from the full rendered image; the sharpness of this specular component is reduced via a normal-based blur operator; the diffuse and specular components are then optionally multiplied by a colour; finally, the intensity and saturation of the resulting specular and diffuse components are adjusted and the final image is obtained by addition of these components (specular + diffuse). These steps are explained in more detail below.

The normal-based blur operator is implemented with a cross bilateral filter (Paris et al., 2009) on the specular component and takes into account the orientation of the surface normals. Such a filter is controlled by a pair of so-called spatial and range weighting functions. The space weight is given by a box kernel of a large static size (160×160 pixels), which merely serves to limit the set of pixels that are considered for blurring. The range weight is given by an exponentiated normal dot product: (**n · n_0_**)^s^, where **n_0_** is the normal at the centre pixel in the kernel, **n** is the normal at a neighbour pixel, and *s* is a shininess parameter. This simple range weighting formula is directly inspired by Phong shading: large shininess values result in a sharp angular filter, whereas small shininess values result in a broad angular filter, which has the effect of blurring specular reflections. In practice, we use a blur parameter *b* in the [0..1] range, which is empirically remapped to shininess using *s = 1 + 1/((b/5)^4^*100+ε)*, where *ε* is a small constant used to avoid division by 0.

Some particular categories of materials such as gold or chocolate require a colorized diffuse or specular component. This is obtained by simply multiplying the image component by a colour: *sCol* for specular, *dCol* for diffuse. In some cases (gold and silver), the diffuse component is entirely discarded, which is equivalent to having *dCol*=(0,0,0) as indicated in Figure 8A. We have made the choice of using bright and saturated colours in all other cases in order to make the parameters of the next step more easily comparable.

For the last intensity and colour adjustment step, each image component is first converted to the HSV colour space. The following manipulations apply either to the specular or diffuse component, with the corresponding parameters either prefixed by ‘s’ or ‘d’. The value channel is manipulated by a combination of gamma exponentiation (controlled by *dGamma* and *sGamma*) and multiplication (by *sBoost* or *dBoost*) using the following simple formula: *Boost*\**v*^*Gamma*^, where *v* stands for value in HSV colour space. The saturation channel is also multiplied (by *sSat* or *dSat*). Both components are converted back to RGB and added together to yield the final image.

We use an additional filter for the specific case of velvet/satin. It is applied to the specular component right after normal-based blurring. The specular component is multiplied by a first mask *m_1_* that is computed via a non-linear mapping of the luminance *l* of the specular component. We use the following formula: *m_1_ = f(0, l_m_/2, l)* if *l<l_m_/2*, and *m_1_=1-f(l_m_/2, l_m_, l)* otherwise, with *f(a,b,x)* a smooth Hermite interpolation between 0 and 1 when *a<x<b*, and *lm* the maximum luminance in the specular component image. The effect of multiplication by *m1* is to darken the core of the brightest highlights, hence producing elongated highlight loops as seen in Supplementary Figures 16 and 17. It is similar in spirit to the sinusoidal modulation of Sawayama and Nishida (Figure 11 in Sawayama & Nishida 2018), except that it is only applied to the specular component and with a different non-linear remapping. We have also found it necessary to multiply the specular component by a second mask *m_2_* to slightly attenuate some of the elongated highlights. This mask is computed using a simple normal dot product: *m_2_=(**n · n_m_**)*, where **n** is the normal at a pixel and **n_m_** is a manually-set direction that is chosen per scene.

#### Manipulation parameters: cue manipulation experiment (Experiment 4)

For each of the four scenes showing either of two objects (bunny and dragon) in either of two lighting environments (campus and kitchen), we have taken as input a material configuration previously classified as ‘Glazed Ceramic’, from which we have produced 12 manipulations for each of the other material categories. There were also two manipulation conditions (see below), yielding a total of 104 stimulus images for the cue manipulation experiment.

The input material was systematically chosen to have a grayscale *Base Color* of 0.1, a *Specular* level of 0.3, a *Specular Tint* of 0, a *Roughness* of 0, and an *Anisotropic* level of 0.

The manipulation parameters are listed in the table below. In the “simple” manipulation condition, all the *Gamma* (i.e., non-linear) parameters were set to 1 and the special velvet filter was discarded. In the “complex” manipulation condition, setting some of the *sGamma* and *dGamma* parameters away from 1 had an impact on the intensity of either the specular or diffuse component; as a result, we also had to adjust the *sBoost* and *dBoost* parameters. We applied the velvet filter only for the velvet manipulation in this condition, using the following **n_m_** directions: (0.57,−0.53,0.63) for Bunny/Campus, (−0.22,−0.48,0.85) for Bunny/Kitchen, (−0.62,0.7,0.35) for Dragon/Campus, and (−0.72,0.57,0.39) for Dragon/Kitchen.

**Table 1:** Filter parameters for the “simple” image manipulation condition

| Category | Blur | sCol | sSat | sBoost | sGamma | dCol | dSat | dBoost | dGamma |
| --- | --- | --- | --- | --- | --- | --- | --- | --- | --- |
| Glazed ceramic | 0 | (1,1,1) | 1 | 1 | 1 | (1,1,1) | 1 | 1 | 1 |
| Brass/Bronze/Copper | 0.45 | (1,0.78,0) | 0.6 | 1.8 | 1 | (1,0.87,0) | 0.25 | 0.75 | 1 |
| Porc./Stone/Chalk | 1 | (1,1,1) | 0 | 0.4 | 1 | (1,1,1) | 0 | 5.5 | 1 |
| Chocolate (liquid) | 0.05 | (1,0.84,0) | 0.4 | 0.5 | 1 | (1,0.75,0) | 0.4 | 1.4 | 1 |
| Chocolate (solid) | 0.5 | (1,0.84,0) | 0.2 | 0.75 | 1 | (1,0.75,0) | 0.4 | 1.4 | 1 |
| Glazed Porcelain | 0 | (1,1,1) | 0 | 0.25 | 1 | (1,1,1) | 0.3 | 6 | 1 |
| Gold metals | 0.15 | (1,0.74,0) | 0.6 | 2.25 | 1 | (0,0,0) | 0.75 | 2.75 | 1 |
| Pearl | 0.525 | (1,1,1) | 0.5 | 1.625 | 1 | (1,1,1) | 0.7 | 3 | 1 |
| Plastic | 0.35 | (1,1,1) | 0.625 | 1 | 1 | (1,1,1) | 0 | 1.75 | 1 |
| Latex/Rubber/Playdoh | 0.75 | (1,1,1) | 1.25 | 0.75 | 1 | (1,1,1) | 1 | 1.5 | 1 |
| Silver metals | 0.125 | (1,1,1) | 0.25 | 1.25 | 1 | (0,0,0) | 1 | 3 | 1 |
| Wax/Soap | 0.35 | (1,1,1) | 0 | 0.25 | 1 | (1,0.84,0) | 0.5 | 6.25 | 1 |
| Velvet/Silk/Fabric | 0.6 | (1,0.95,0) | 0.33 | 1.25 | 1 | (1,0.82,0) | 0.6 | 3.75 | 1 |

**Table 2:** Filter parameters for the “complex” image manipulation condition. Modified parameters compared to Table 1 are shown in bold.

| Category | Blur | sCol | sSat | sBoost | sGamma | dCol | dSat | dBoost | dGamma |
| --- | --- | --- | --- | --- | --- | --- | --- | --- | --- |
| Glazed ceramic | 0 | (1,1,1) | 1 | 1 | 1 | (1,1,1) | 1 | 1 | 1 |
| Brass/Bronze/Copper | 0.45 | (1,0.78,0) | 0.6 | **3** | **1.4** | (1,0.87,0) | 0.25 | **1.25** | **1.2** |
| Porc./Stone/Chalk | 1 | (1,1,1) | 0 | 0.4 | 1 | (1,1,1) | 0 | 5.5 | 1 |
| Chocolate (liquid) | 0.05 | (1,0.84,0) | 0.4 | 0.5 | 1 | (1,0.75,0) | 0.4 | 1.4 | 1 |
| Chocolate (solid) | 0.5 | (1,0.84,0) | 0.2 | **1.6** | **1.5** | (1,0.75,0) | 0.4 | 1.4 | 1 |
| Glazed Porcelain | 0 | (1,1,1) | 0 | **1** | **1.8** | (1,1,1) | 0.3 | **2.3** | **0.5** |
| Gold metals | 0.15 | (1,0.74,0) | 0.6 | **1.625** | **0.42** | (0,0,0) | 0.75 | 0 | 1 |
| Pearl | 0.525 | (1,1,1) | 0.5 | **1.1** | **0.6** | (1,1,1) | 0.7 | 2.37 | 1 |
| Plastic | 0.35 | (1,1,1) | 0.5 | **0.8** | **1.3** | (1,1,1) | 0 | **1.5** | **0.9** |
| Latex/Rubber/Playdoh | 0.75 | (1,1,1) | 1.25 | 0.75 | 1 | (1,1,1) | 1 | 1.5 | 1 |
| Silver metals | 0.125 | (1,1,1) | 0.25 | **1.25** | **0.3** | (0,0,0) | 1 | 0 | 1 |
| Wax/Soap | 0.35 | (1,1,1) | 0 | **0.13** | **0.7** | (1,0.84,0) | 0.5 | **2.2** | **0.5** |
| Velvet/Silk/Fabric | 0.6 | (1,0.95,0) | 0.33 | 2.9 | 1 | (1,0.82,0) | 0.6 | 3.75 | 1 |

### Procedure

#### Participants and stimulus presentation

Participants were students at Justus Liebig University Giessen in Germany and were compensated 8 euros per hour. The experiments followed the guidelines set forth by the Declaration of Helsinki.

In the free naming task, stimuli were projected onto a white wall in a classroom. Thick black cloth was used to block light from the windows, so that the only source of light came from the projected image. For all other experiments the stimuli were presented on a Sony OLED monitor running at a refresh rate of 120 Hz with a resolution of 1920 × 1080 pixels controlled by a Dell computer running Windows 10. Stimuli were viewed in a dark room at a viewing distance of approximately 60cm. The only source of light was the monitor that displayed the stimuli.

Stimulus presentation and data collection were controlled by a MATLAB script (release 2018b, Mathworks, Natick, MA) using the Psychophysics Toolbox (Brainard, 1997).

#### Free-naming task (Experiment 1)

Fifteen participants completed free naming experiment. All participants gathered in a classroom and completed the task at the same time. They viewed stimuli one at a time and were asked to classify the material of each stimulus, with no restrictions. They were provided with sheets of paper with space for each trial number to write down their answers. The experimenter controlled stimulus presentation with a keyboard press. Each trial was presented for as long as it took for all participants to finish writing down their responses. A blank screen was shown between each trial for one second. The experiment took approximately three hours to complete, including breaks.

The instructions, written on a sheet of paper in both English and German, were as follows:

> You will be shown 270 images, and your task is to write down your impressions of the material of each object, i.e. what does it look like it is made of? Below are some suggestions of materials that might help prompt you. Your answers might not include, nor are they limited to, the suggestions below. There are no right or wrong answers and you can respond however you like. You can be as specific (e.g. aluminium foil, polyethylene), or general (e.g. metal, plastic) as you like. You can also write down more than one material (e.g. “looks most like glazed ceramic but could also be plastic”), or even say that it doesn’t look like anything you know. If you can’t remember the name of a material, you can write e.g. “the stuff that X is made out of”.

The following examples were provided below the instructions:

- Metal, e.g. silver, gold, steel, iron, aluminium, chrome, foil
- Textiles, e.g. velvet, silk, leather
- Plastic, e.g. PVC, nylon, acrylic, polyethylene, Styrofoam
- Ceramic, e.g. porcelain, china
- Minerals, e.g. stone, concrete, rock
- Coatings, e.g. glazed, painted, enamel, PTFE/Teflon
- Other: soap, wax, chalk, pearl, composite materials

#### 18-AFC task (Experiment 2)

Eighty native-level German speakers participated in the 18-AFC experiment. The 924 stimuli were randomly split into 4 sessions, so that each participant categorised a quarter of the stimuli (20 participants per stimulus). The experiment was self-paced with no time constraints, and for most participants the experiment lasted approximately 1-1.5 hours.

On each trial, observers were presented with an object in the centre of the screen (29° visual angle) with 18 categories displayed along the sides of the stimulus (Figure 4A). They were asked to choose the category that best applied to that stimulus. If they were unhappy with their choice, they could change the confidence rating at the bottom right of the screen. Before the experiment, participants were asked to read carefully through the list of materials and were shown several examples of the stimuli to be presented. The purpose of this was so observers got a sense of the range of materials in the experiment. Observers were restricted to choosing only one category for each stimulus, and were given the following instructions verbally by the experimenter:

> Use the mouse to click on the category that best describes the material of each object. When you are satisfied with your choice press the space bar to proceed to the next trial. In the case of categories with multiple items (e.g. velvet/silk/fabric), the perceived material only needs to apply to one, not all, the categories. There are no right or wrong answers, as the experiment is about the perception of materials. Not all categories will have an equal number of stimuli – you may choose one category more or less than others (or not at all). If you are not satisfied or confident with your choice, change the confidence rating at the bottom of the screen. This should be used in the case you feel that none of the available categories fit; if you think more than one category applies then just choose the most suitable option.

#### Gloss rating task (Experiment 3)

Twenty-two native-level German and English speakers participated in the gloss ratings experiment. One participant was excluded because they did not understand the task instructions, resulting in 21 participants. The experiment was self-paced with no time constraints, and for most participants the experiment lasted approximately 1 hour.

Before the experiment, participants were shown real world examples of glossy objects (some of which are shown in Figure 1A). As they were shown these images, they were given the following instructions:

> Many objects and materials in our environment are glossy or shiny. Glossy things have specular reflections, and different materials look glossier/shinier than others. You will be shown different objects and asked to rate how glossy each object looks. This experiment is about visual perception so there is no right or wrong answer – just rate how glossy each object looks to you.

Participants were shown many example trials before starting the experiment so that they could experience the full range of stimuli and calibrate their ratings to the range of gloss levels in the experiment. On each trial, observers were presented with an object in the centre of the screen (29° visual angle). The task was to rate how glossy the object was by moving the mouse vertically to adjust the level of a bar on the right side of the screen. They were told to look at many points on the object before making their decision. The starting level of the rating bar was randomly set on each trial.

#### Cue manipulation experiment (Experiment 4)

Twenty-two native-level German speakers participated in the cue manipulation experiment. Each participant categorised all 104 stimuli, and the task was identical to the 18-AFC task in Experiment 2. The experiment was self-paced with no time constraints, and for most participants the experiment lasted approximately 45 minutes.

## ANALYSES

### Free-naming experiment (Experiment 1)

A student assistant went through participants’ responses of the free-naming task and extracted all category terms (nouns) and descriptors (e.g., adjectives describing fillings, coatings, finishes, and states), translating them to English. Each participant often used multiple terms per stimulus, all of which were recorded as separate entries. Similar terms (like foil/metal foil/aluminium foil, or pearl/pearlescent/mother of pearl/bath pearl) were combined into a single category. Some terms like “Kunststoff” and “Plastik” were considered duplicates because they translated to a single term (plastic) in English.

### 18-AFC experiment (Experiment 2)

#### Reduced set of category terms

Two of the authors (AS and KD) worked together to reduce the set of category terms that were generated in the free-naming task (the 29 category terms that at least 5 out of 15 participants used; see Supplementary Figure 2). Note that this was an arbitrary cut off with the aim of being quite inclusive while at the same time maintaining some consensus among participants. Materials that were visually or semantically similar were combined (specifically: fabrics, including velvet and silk; unglazed porcelain, stone, and chalk; latex and rubber; wax and soap; brass, bronze, and copper; iron and chrome; and silver and aluminium). This was guided by correlations between the categories (based on the number of participants that used each term for each stimulus; Supplementary Figure 4). The superordinate category “metal” was not included separately. In the free-naming task, non-category terms were often required to distinguish particular coatings (glazed/varnished), finishes (polished), or states (liquid/melting; see Supplementary Figure 2). For the 18-AFC task, we separated categories where these terms applied; for example, we included both glazed porcelain and unglazed porcelain, both liquid and solid chocolate, and also included a “covered in wet paint” category. Thus, the final set of 18 category terms (Supplementary Figure 5) that participants could choose from in the 18-AFC task was as inclusive as possible while not being too large to overwhelm participants.

#### Factor analysis

The correlations between different categories in terms of their stimulus profiles (Supplementary Figure 6) indicate that some category terms were not independent. This suggests the existence of underlying common dimensions; that is, participants used the same underlying criteria for different category terms. An exploratory factor analysis was performed on the stimulus profiles from Figure 3B, which allowed us to explain some of this covariation and reveal the underlying category space of our stimulus set. Figure 4A (red squares) shows that there is a steady increase in the common variance explained by each additional factor. The amount of additional variance explained by each factor did not drop off after a certain number of factors, indicated by the absence of a plateaux in this plot. Therefore, we extracted 12 factors (the upper limit based on degrees of freedom), which where interpretable and explained over 80% of the common variance between stimulus profiles. This is similar to or greater than the amount of variance explained by the factors/components retained in other material perception studies that have used factor analysis or principal components analysis (see Schmid & Doerschner, 2018, for a discussion of these studies). The factors were interpretable (Figure 4B) and were labelled based on the original categories that loaded most strongly onto each factor (Figure 4C). Factor scores were calculated using a weighted least-squares estimate (also known as the “Bartlett” method). For comparison, Supplementary Figure 7 shows the results of a principal components analysis that retained all dimensions in addition to the results of other factor solutions, whose dimensions overlap with those here.

#### Visual feature (cue) measurements

Calculations for most of these cues relied on specular highlight coverage maps. For glossy surfaces with uniform reflectance properties (like the stimuli used in the present study), specular reflections cover the entire surface. However, for low-gloss objects we often only see specular highlights, or “bright” reflections, which are usually reflections of direct light sources like the sun, or a lamp. For very shiny surfaces (like metal) and in some lighting conditions we also see lowlights, or “dark” reflections (Kim et al., 2012), which are reflections of indirect light coming from other surfaces in the scene. We chose to define *coverage* as the amount of the surface covered in specular highlights, excluding lowlights, which is consistent with previous literature (Marlow et al., 2012; Di Cicco, Wijntjes, & Pont, 2019). Marlow and colleagues measured *coverage*, *contrast*, and *sharpness* of specular highlights using perceptual judgments from participants, due to the difficulty in segmenting the specular component of an image from other sources of image structure (such as diffuse reflectance and shading). An acknowledged concern of this approach is that it uses one perceptual output (perceived coverage, contrast, and sharpness) to match another (perceived gloss). It is unclear what image structure observers use to judge each cue, and participants might conflate their judgments of the visual cues with each other and with perceived gloss (Marlow & Anderson, 2013; see also van Assen, Barla, & Fleming, 2018, for a similar use of this method). Therefore, we wanted to develop objective measures of specular reflection cues. Currently there is no established method for segmenting specular reflections and diffuse shading from a single image that is robust across different contexts (e.g., changes in surface albedo, shape, and illumination conditions). To help with this segmentation we rendered additional images that isolated specular and diffuse components.

##### Extra rendered images

For each stimulus, a purely specular image was rendered, which had the same specular reflectance as the original (full rendered) image but with diffuse component turned off. Two purely diffuse images were rendered, which we call “diffuse image 1” (used to calculate *coverage*) and “diffuse image 2” (used to calculate *sharpness* and *contrast*; first column in Supplementary Figure 9). Kim et al. (2012) separated specular highlights and lowlights by subtracting an image of a rendered glossy surface (with a specular and diffuse component) from an “equivalent” fully diffuse image with the same total reflectance as the glossy surface. Kim et al. used the Ward model (Ward, 1994) where the total reflectance could be easily matched between glossy and diffuse renderings because the glossy surface reflected the same amount of incident light at all viewing angles. Since the Principled BSDF simulates the Fresnel effect (whereby specular reflectance increases at grazing viewing angles, depending on the index of refraction), the diffuse (1) renderings were matched to the purely specular renderings in total reflectance at facing surface normal (i.e., surface orientations that face the camera). The second diffuse image (diffuse image 2) was created by rendering only the diffuse component of the original (full rendered) stimulus, with the specular component turned off.

##### Segmentation

Specular reflections were segmented into specular highlights and lowlights by subtracting the diffuse image 1 from the purely specular image, which resulted in a subtracted image. This subtracted image was thresholded to serve as a coverage mask (second column in Supplementary Figure 9). For best results for our stimulus set, the threshold was set to one third of the maximum diffuse shading for each stimulus. Pixels above this threshold were considered highlights, and the remaining pixels were considered lowlights. For calculations of sharpness and contrast, specular reflections were segmented from diffuse shading by subtracting the diffuse image 2 from the original “full” stimulus (second column in Supplementary Figure 9).

##### Cues

*Coverage* was defined as the proportion of object pixels that were calculated to be specular highlights (excluding lowlights). *Contrast* was the sum of root-mean-squared (RMS) contrast of extracted specular reflections at different spatial frequency bandpasses. *Sharpness* of extracted specular reflections was calculated for each pixel within the highlight regions using a measure of local phase coherence (Hassen, Wang, & Salama, 2013), then these values were averaged. *Highlight saturation* and *lowlight saturation* were calculated as the average colour saturation of pixels within the highlight region, and outside of the highlight region (which we call the lowlight region), respectively. *Lowlight value* was calculated as the average value of pixels in the lowlight region. A root transformation on sharpness and contrast was applied to linearize the relationship of these cues to gloss ratings.

### Gloss ratings experiment (Experiment 3)

#### Correlations

Intersubject correlations for gloss ratings were calculated using Pearson correlation, and the median was taken. Pearson correlations used for analyses in Figure 5D and F were Fisher-transformed.

#### Linear regression

A Linear regression was performed on 920 stimuli predicting mean gloss ratings from the three gloss cues (coverage, sharpness, contrast). Only stimuli that were allocated a dimension in the factor analysis (from Experiment 2) were included in this analysis (four stimuli loaded negatively onto dimensions, but not enough to make a new dimension for our stimulus set).

### Cue manipulation experiment (Experiment 4)

The same image cue measurements were used for the rendered stimuli (from Experiment 2) and the manipulated images (from Experiment 4), so that the measurements would be comparable (see Supplemental Figure 14). However, slight modifications had to be made. For the rendered stimuli (Experiment 2), the segmentation between highlights and lowlights (described in the previous section) relied on two additional rendered images: a rendered specular image and a rendered diffuse image (1) with the same total reflectance (Supplementary Figure 9). For Experiment 4, the image cue manipulations are all applied to the same “Glazed Ceramic” material (see Methods). We reused the corresponding rendered specular and diffuse images from the stimulus set from Experiment 2, with one additional step: the specular image was further modified by the normal-based blur filter to account for the change in highlight coverage induced by a change in sharpness. Apart from this, the rest of the image cue measurement routines remained unchanged.

## ACKNOWLEDGEMENTS

A.C.S. and K.D. are supported by a Sofja Kovalevskaja Award endowed by the German Federal Ministry of Education, awarded to K.D. P.B. is supported by ANR project VIDA (ANR-17-CE23-0017). We thank Marios Panayi, Chris Baker, Filipp Schmidt, and Karl Gegenfurtner for helpful feedback on earlier versions of this manuscript.

