## Supplemental Figures for "Material category of visual objects computed from specular image structure"

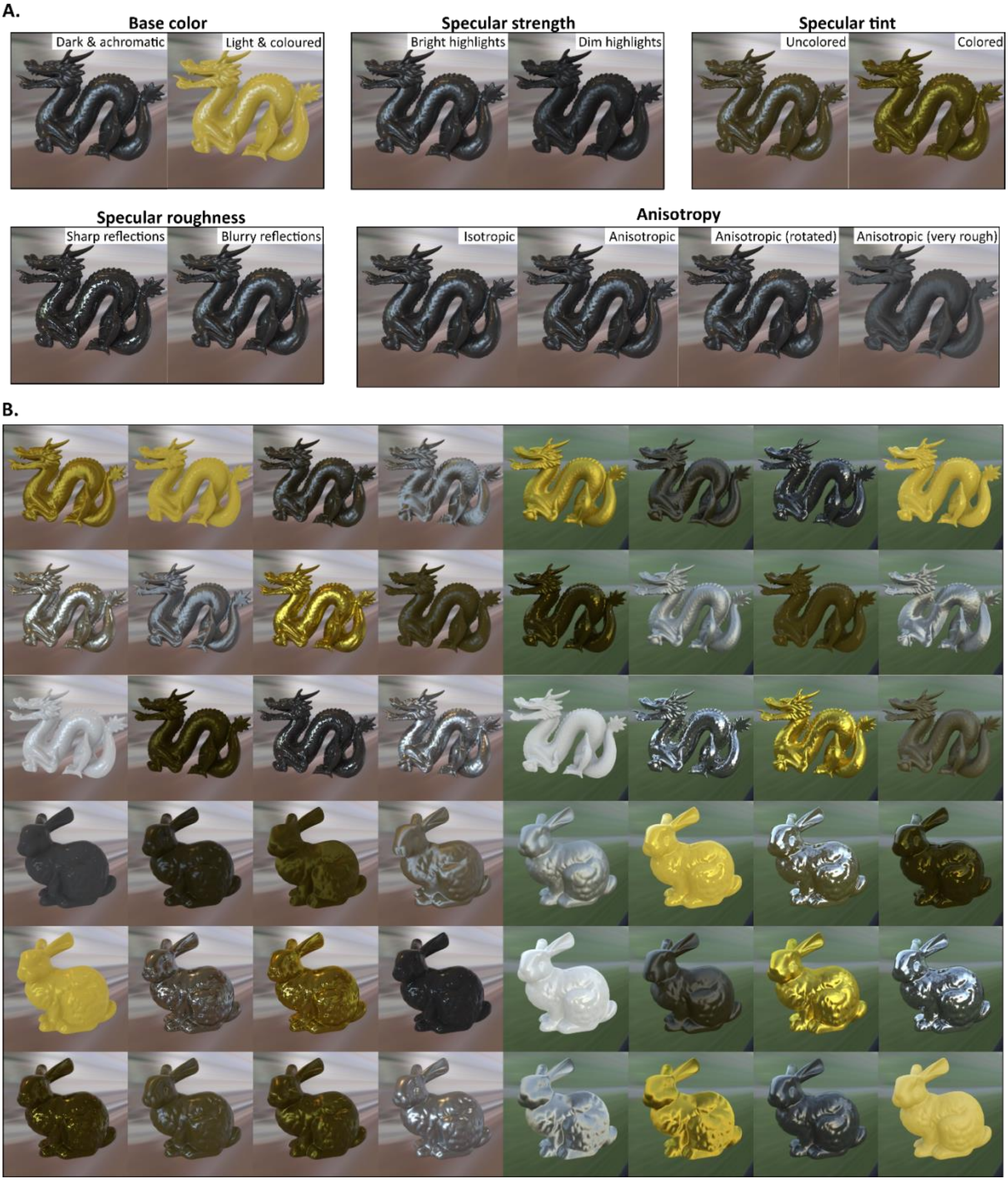


**Supplementary Figure 1.** Computer-rendered stimuli in the experiments were complex 3D shapes embedded in natural illumination fields. **A.** The visual effects of manipulating different rendering parameters. **B.** A sample of the stimuli used in the 18-AFC and gloss rating experiments (Experiments 2 and 3). Only reflectance parameters were manipulated and yet we found that these variations yielded many different perceived material classes beyond those defined by the reflectance function used.

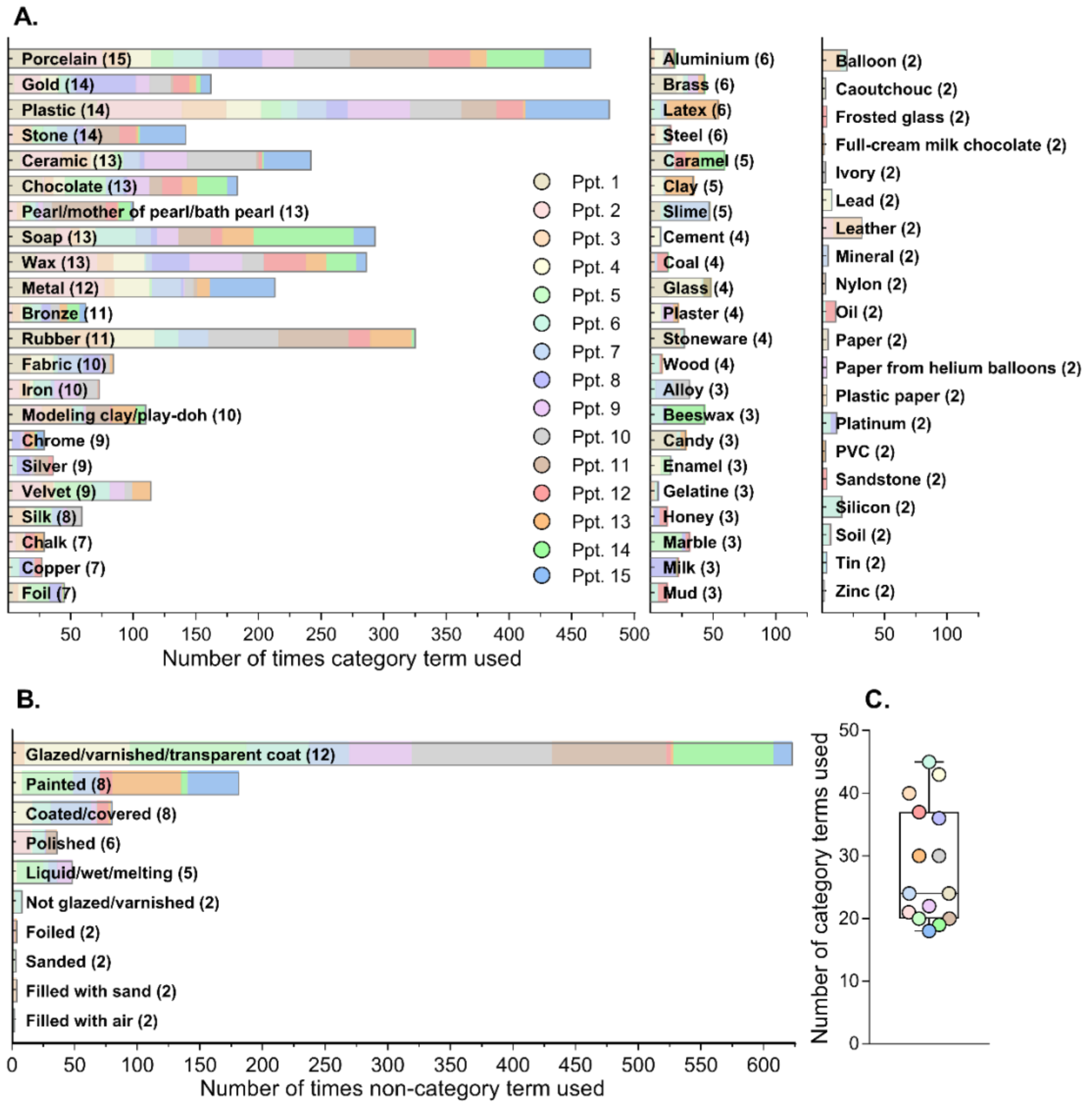


**Supplementary Figure 2.** Results of the free-naming experiment (Experiment 1, n=15). After processing for duplicates and similar terminology, 209 terms were used to describe the materials, 121 of which were *category terms* (i.e., nouns like porcelain, gold, plastic). There were 17 *non-category terms* that included descriptions of fillings (e.g., filled with sand), coatings (e.g., glazed, varnished, painted, covered), finishes (e.g., polished, sanded), and states (e.g., liquid, wet, melting). Participants also used a total of 71 *other adjectives* to describe the stimuli (e.g., dark, yellow, glossy, matte). These other adjectives were excluded from analyses altogether but can be found in Supplementary Figure 3. The bar plots show the number of times each term was used by each participant, for the 64 category terms **(A)** and 10 non-category terms **(B)** that were used by at least two participants. The numbers in brackets correspond to the number of participants (out of 15) that used each term. Not shown are the 57 category terms and 7 non-category terms that were used by only one participant, and the 71 *other adjectives*. Importantly, the use of each term was distributed quite well among participants, i.e., category labels did not come from the same few participants. The box plot in **(C)** supports this by showing that each participant used many category terms (range = 18-45, median = 24).

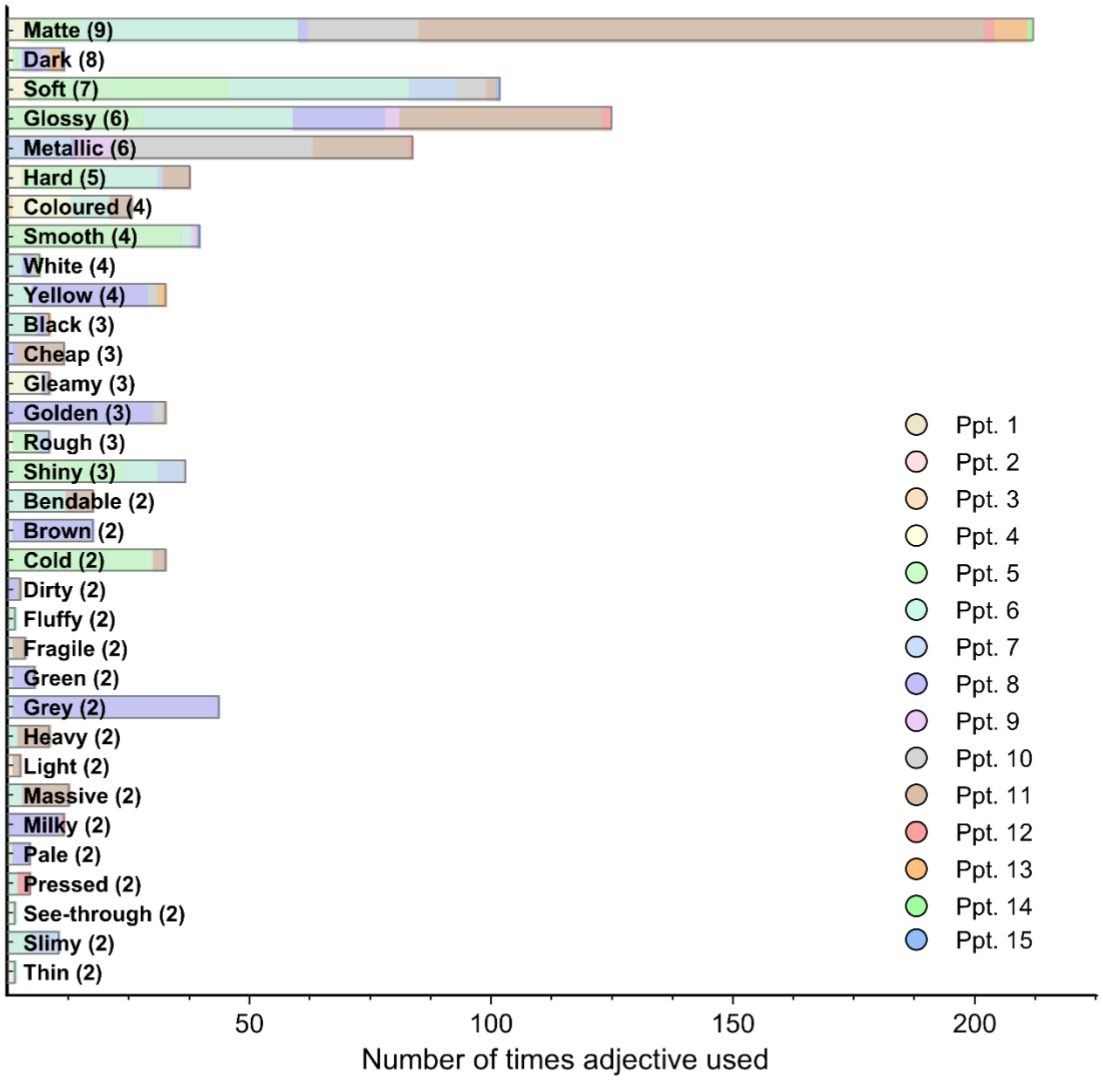


**Supplementary Figure 3.** Additional results from the free-naming experiment (Experiment 1), showing the number of times each *other adjective* term was used. Only adjectives that were used by at least two participants are shown.

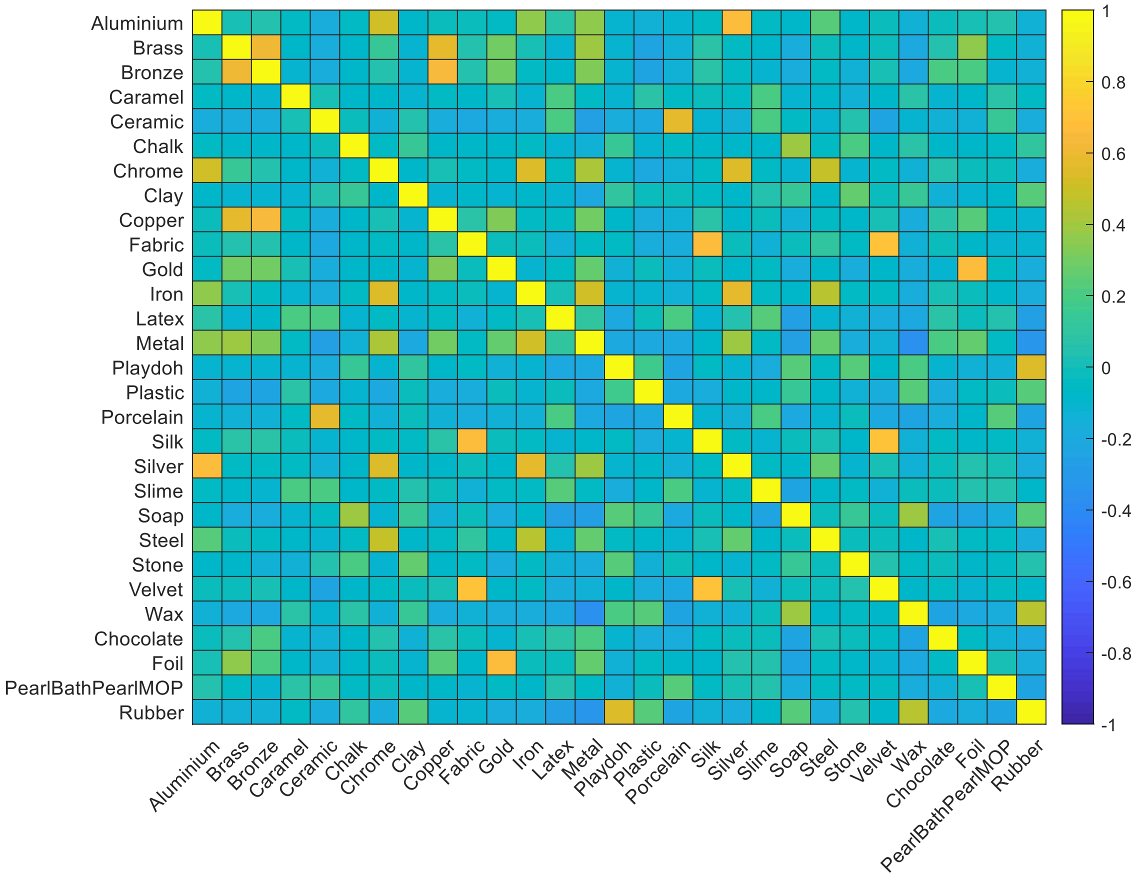


**Supplementary Figure 4.** Heat plot showing Pearson correlations between pairs of categories from the free-naming task (Experiment 1), calculated from the number of participants that used each term for each stimulus. These correlations were used to guide the merging of visually/semantically similar category terms for the 18-AFC task (Experiment 2):

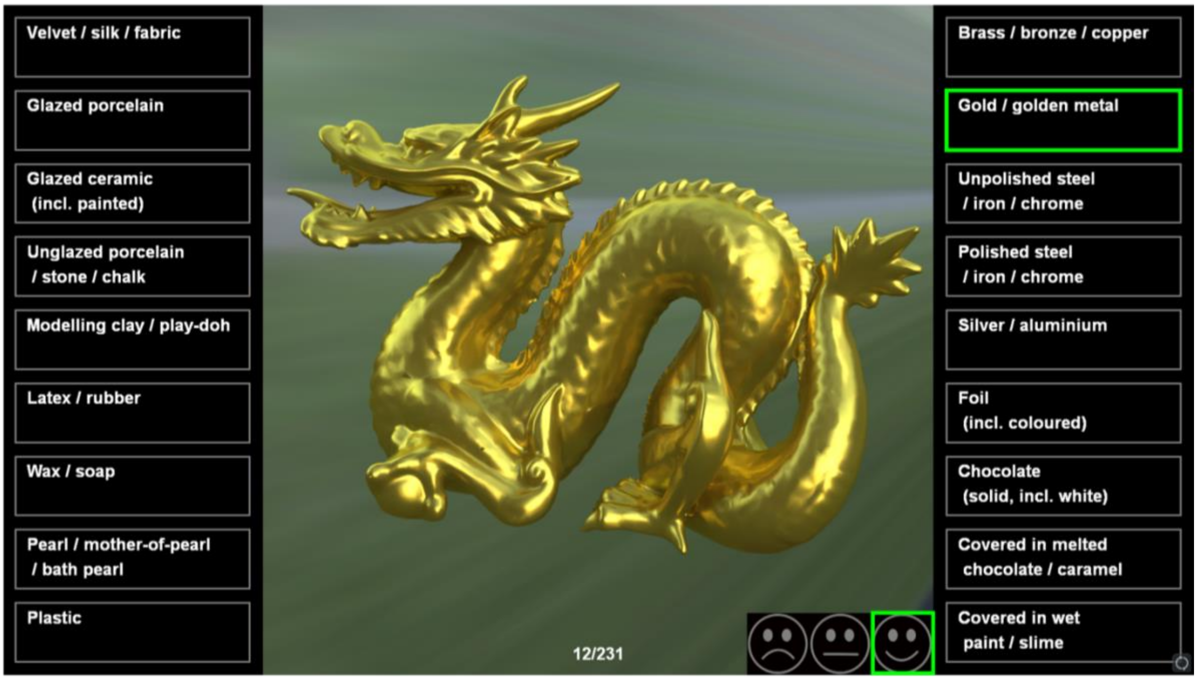


**Supplementary Figure 5.** A screenshot of an example trial from the 18-AFC experiment (Experiment 2). Observers were presented one stimulus at a time and were asked to choose the category that best applied to each stimulus. If they were not perfectly happy with their category choice (e.g., if the category they were looking for was not an option), they could adjust their confidence rating at the bottom right of the screen accordingly. At the bottom centre of the screen was a trial counter indicating the current trial and total number of trials in that block. Note that the categories were presented in German in the actual experiment. The German translations are as follows, in the order shown in the figure (1-9 displayed on the left; 10-18 displayed on the right):

1) Samt / Seide / Stoff
2) Glasiertes Porzellan
3) Glasiertes Steingut / Keramik
4) Unglasiertes Porzellan / Stein / Kreide
5) Knete / Modelliermasse
6) Latex / Gummi
7) Wachs / Seife
8) Perle / Perlmutt / Badeperlen
9) Plastik / Kunststoff
10) Messing / Bronze / Kupfer
11) Gold / goldenes Metall
12) Unpolierter(s) Stahl / Eisen / Chrom
13) Hochpolierter(s) Stahl / Eisen / Chrom
14) Silber / Aluminium
15) Folien (auch farbige)
16) Schokolade (fest, auch weisse)
17) mit flüssiger(m) Schokolade / Karamel überzogen
18) mit flüssiger(m) Farbe / Schleim bedeckt

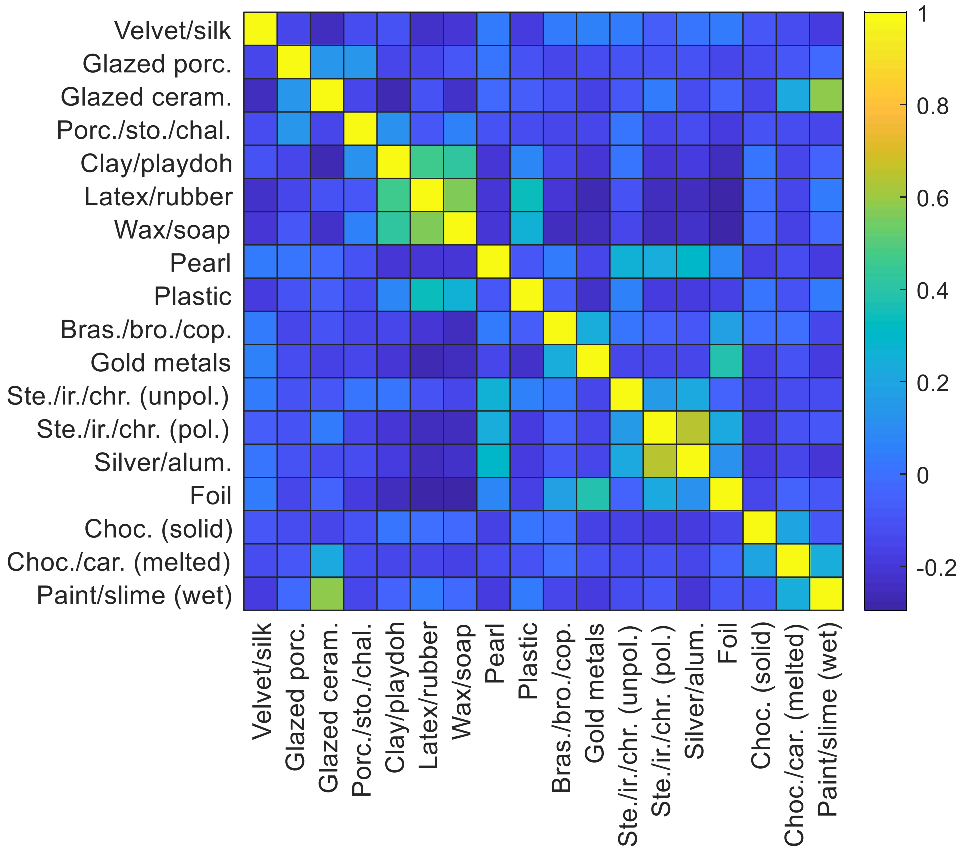


**Supplementary Figure 6.** Heat plot showing Pearson correlations between stimulus profiles (confidence rating sums) for each pair of categories from the 18-AFC experiment (Experiment 2).

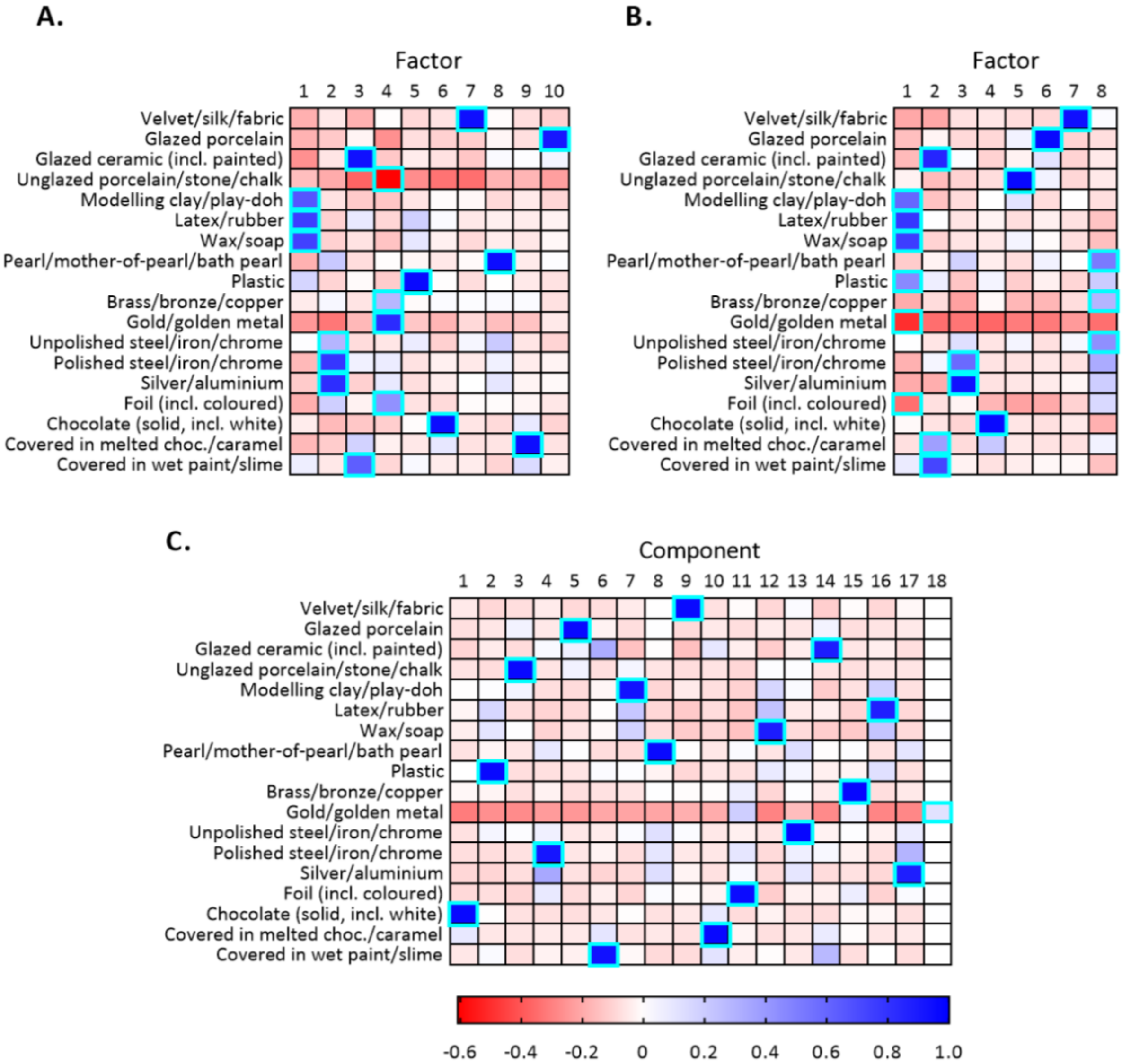


**Supplementary Figure 7.** Heat plots of loadings for each category from the 18-AFC task when different numbers of dimensions are retained. The highlighted cells show the dimension onto which each category most strongly loaded. The emergent dimensions from each analysis are comparable to the 12-factor solution in Figure 4 in terms of their interpretation. **A.** 10-factor solution: (1) Rubber-like; (2) Uncolored metals; (3) Ceramics; (4) Gold metals; (5) Plastic; (6) Solid chocolate; (7) Velvety/silky; (8) Pearlescent; (9) Melted chocolate, (10) Glazed porcelain; (11) Unglazed porcelain **B.** 8-factor solution: (1) Plastic/rubber-like; (2) Ceramics; (3) Uncolored metals; (4) Solid chocolate; (5) Unglazed porcelain; (5) Glazed porcelain; (6) Velvety/silky; (7) Pearlescent/unpolished metals. **C.** Principal components analysis solution retaining all categories.

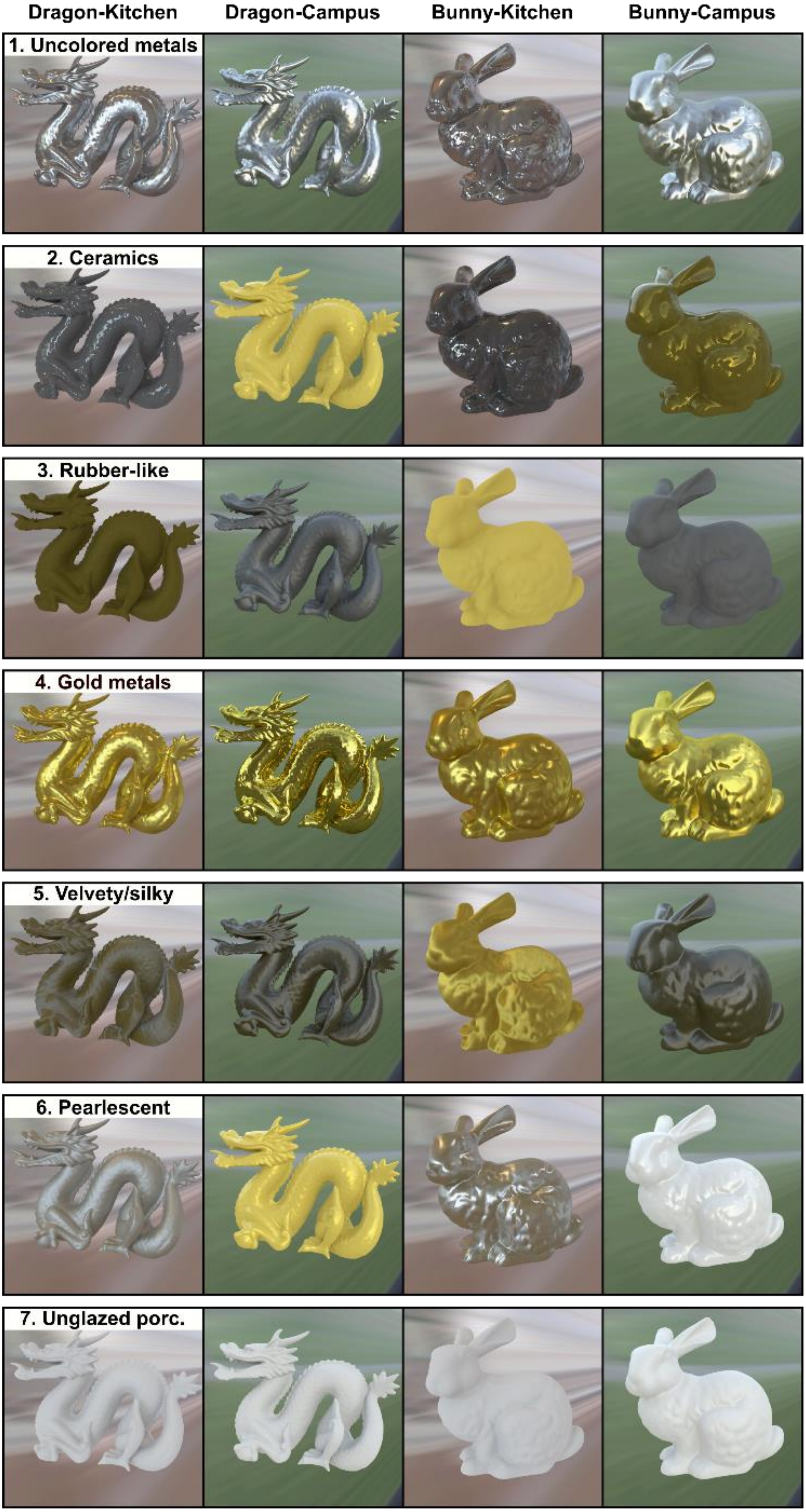
Dragon-Kitchen
Dragon-Campus
Bunny-Kitchen
Bunny-Campus
1. Uncolored metals
2. Ceramics
3. Rubber-like
4. Gold metals
5. Velvety/silky
6. Pearlescent
7. Unglazed porc.

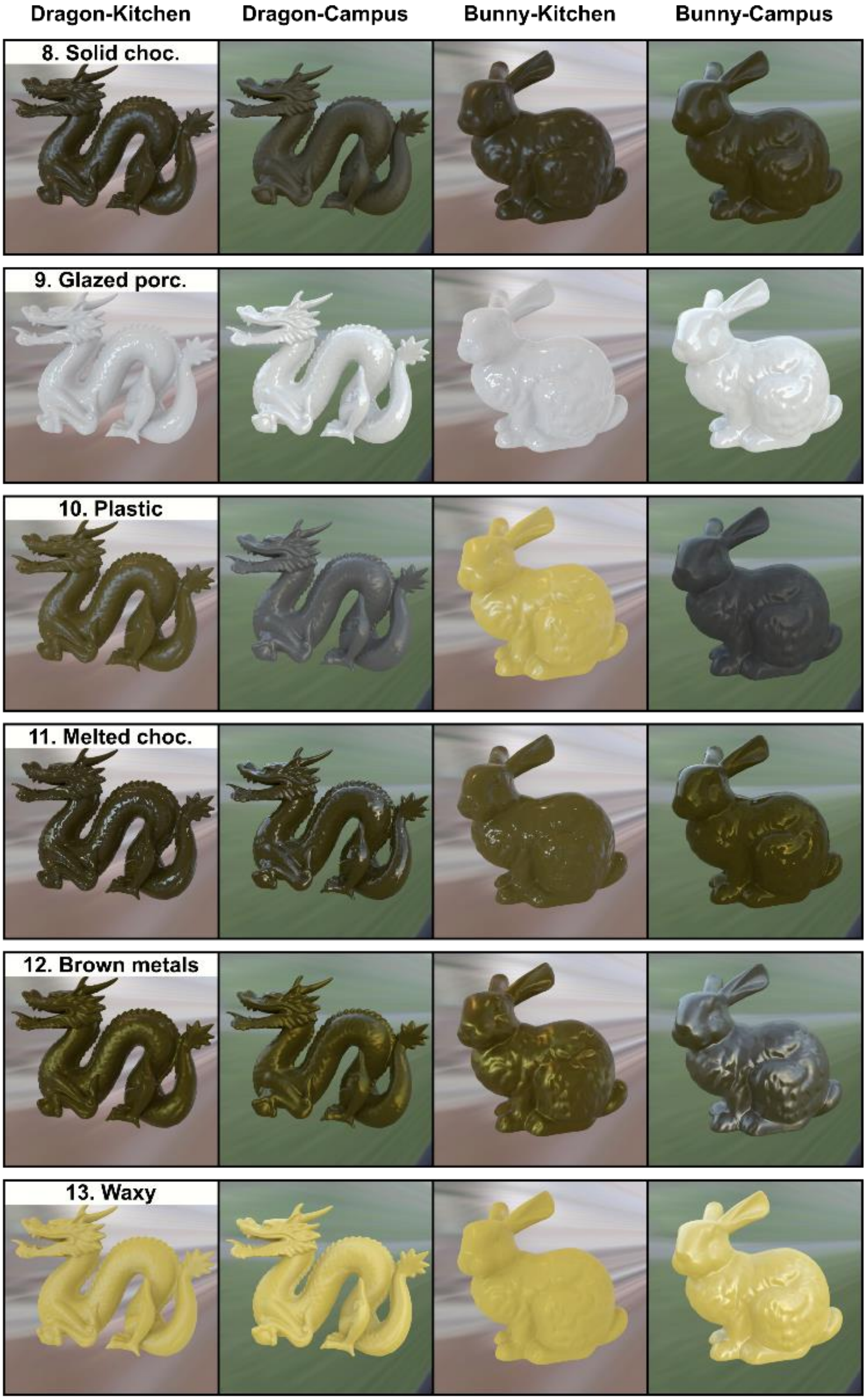


**Supplementary Figure 8.** More example stimuli from each emergent dimension from the factor analysis (Figure 4), for each shape and light field condition.

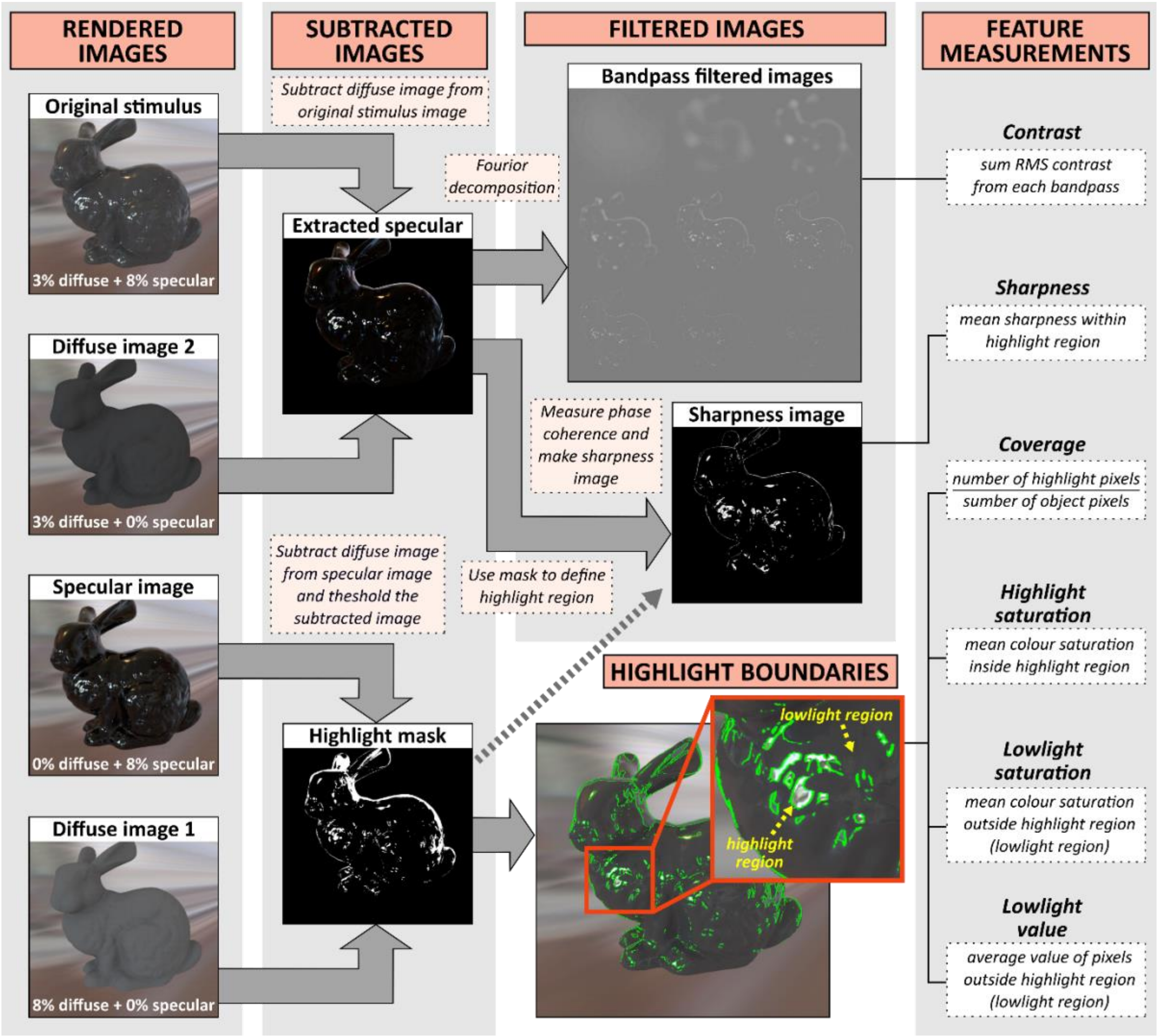


**Supplementary Figure 9.** Calculation of the six specular reflection image cues. See Analyses for details. **First column:** To help with segmenting the image structure caused by specular reflections and diffuse shading, we rendered additional images that isolated specular and diffuse components. **Second column:** We extracted both specular highlights (bright reflections; "Highlight mask" image) and specular reflections, which include both specular highlights and lowlights (i.e., both bright and dim reflections; "Extracted specular" image). **Third column:** The bottom image shows the results of the specular highlight segmentation, with the green outline showing the boundary between highlights and lowlights. The top images show filtered images that are outputted as intermediate steps in the calculation of contrast and sharpness image cues. **Last column:** *Coverage* was defined as the proportion of object pixels that were calculated to be specular highlights (excluding lowlights), based on a threshold above the maximum diffuse shading. *Contrast* was the sum of root-mean-squared (RMS) contrast of extracted specular reflections at different spatial frequency bandpasses. *Sharpness* of extracted specular reflections was calculated for each pixel within the highlight regions using a measure of local phase coherence (Hassen, Wang, & Salama, 2013), then these values were averaged. *Highlight saturation* and *lowlight saturation* were calculated as the average colour saturation of pixels within the highlight region, and outside of the highlight region (which we call the lowlight region), respectively. *Lowlight value* was calculated as the average value (or lightness) of pixels in the lowlight region.

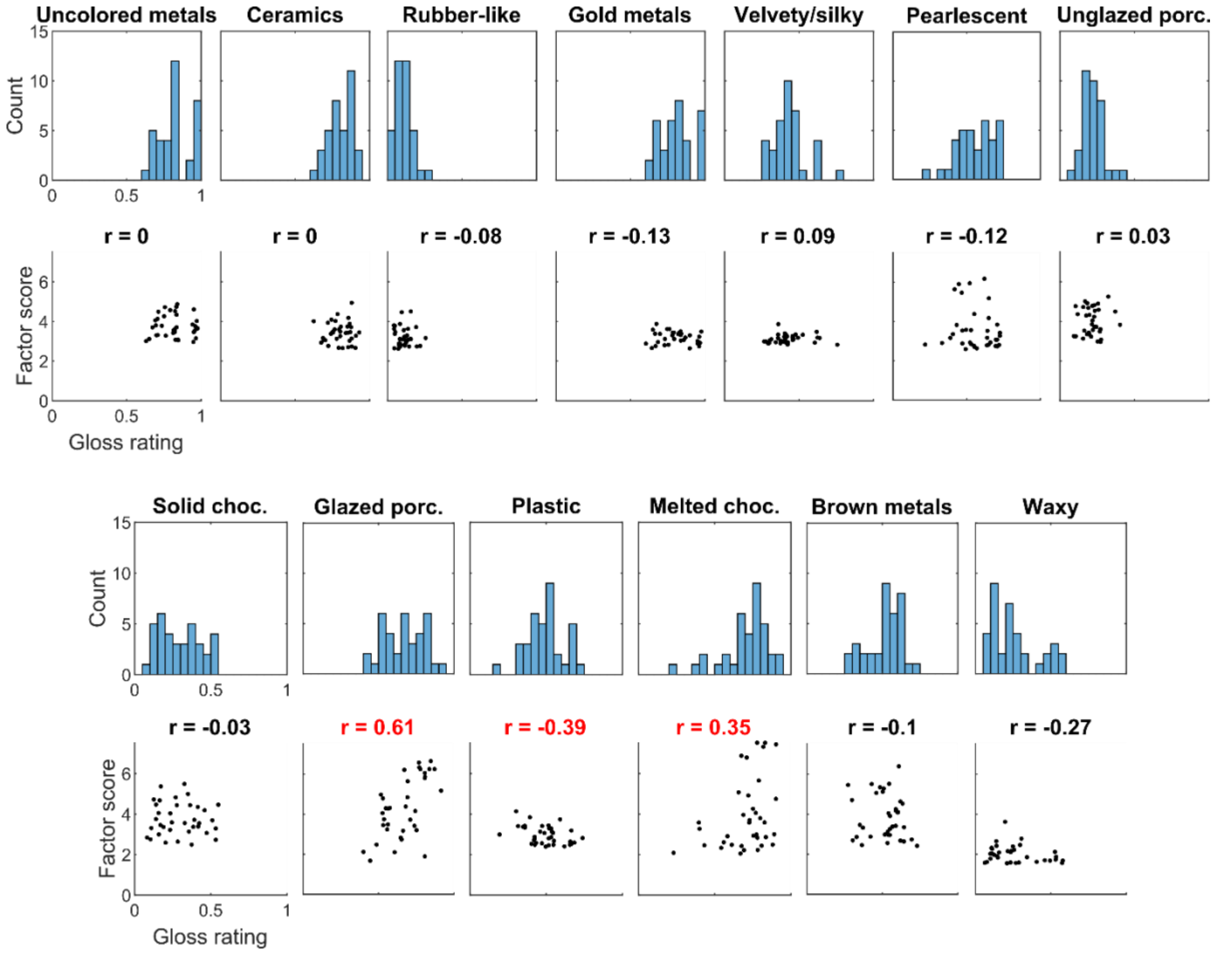


**Supplementary Figure 10.** Histograms and corresponding scatter plots show gloss ratings and correlations between gloss ratings and factor scores (12 factor solution) for the top loading stimuli (cutoff=36). For this more conservative analysis it is apparent that, overall, stimuli from the same material class exhibited a wide distribution of gloss levels, as we saw it in the more inclusive plots in Figure 5.

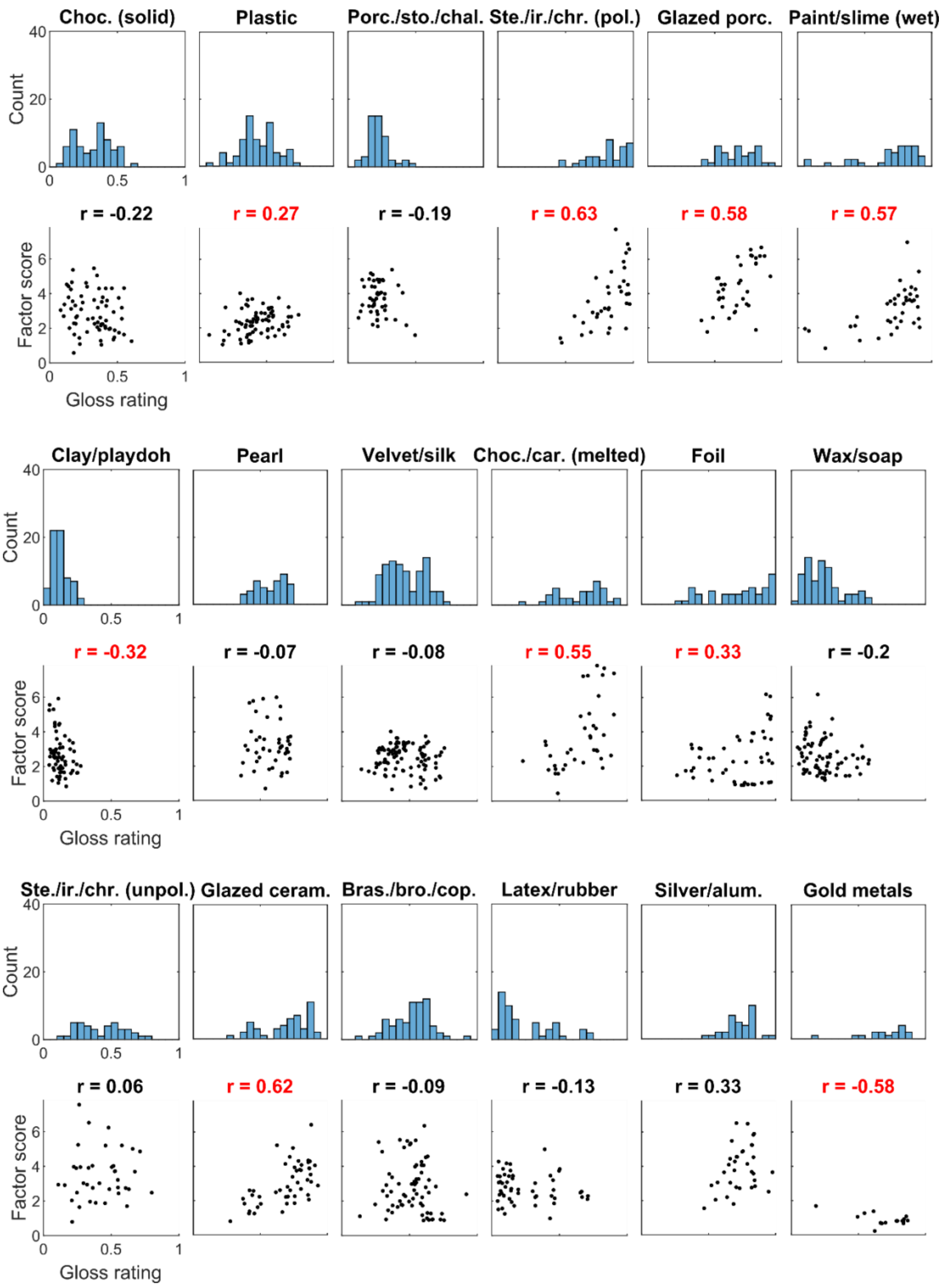


**Supplementary Figure 11.** Histograms and corresponding scatter plots show gloss ratings and correlations between gloss ratings and factor scores (18 factor solution) for all stimuli. The general pattern that we saw in Figure 5 and Supplementary Figure 10 persists, i.e., that stimuli from the same material class exhibit a wide distribution of gloss levels.

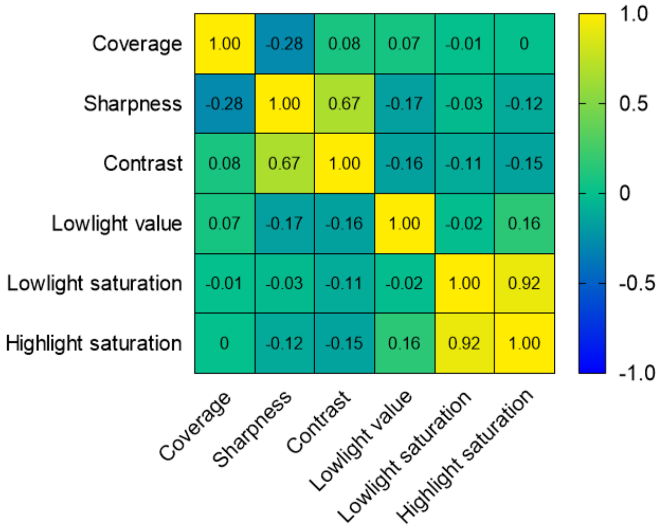


**Supplementary Figure 12.** Heat plot showing Pearson correlations between measured visual features across all stimuli.

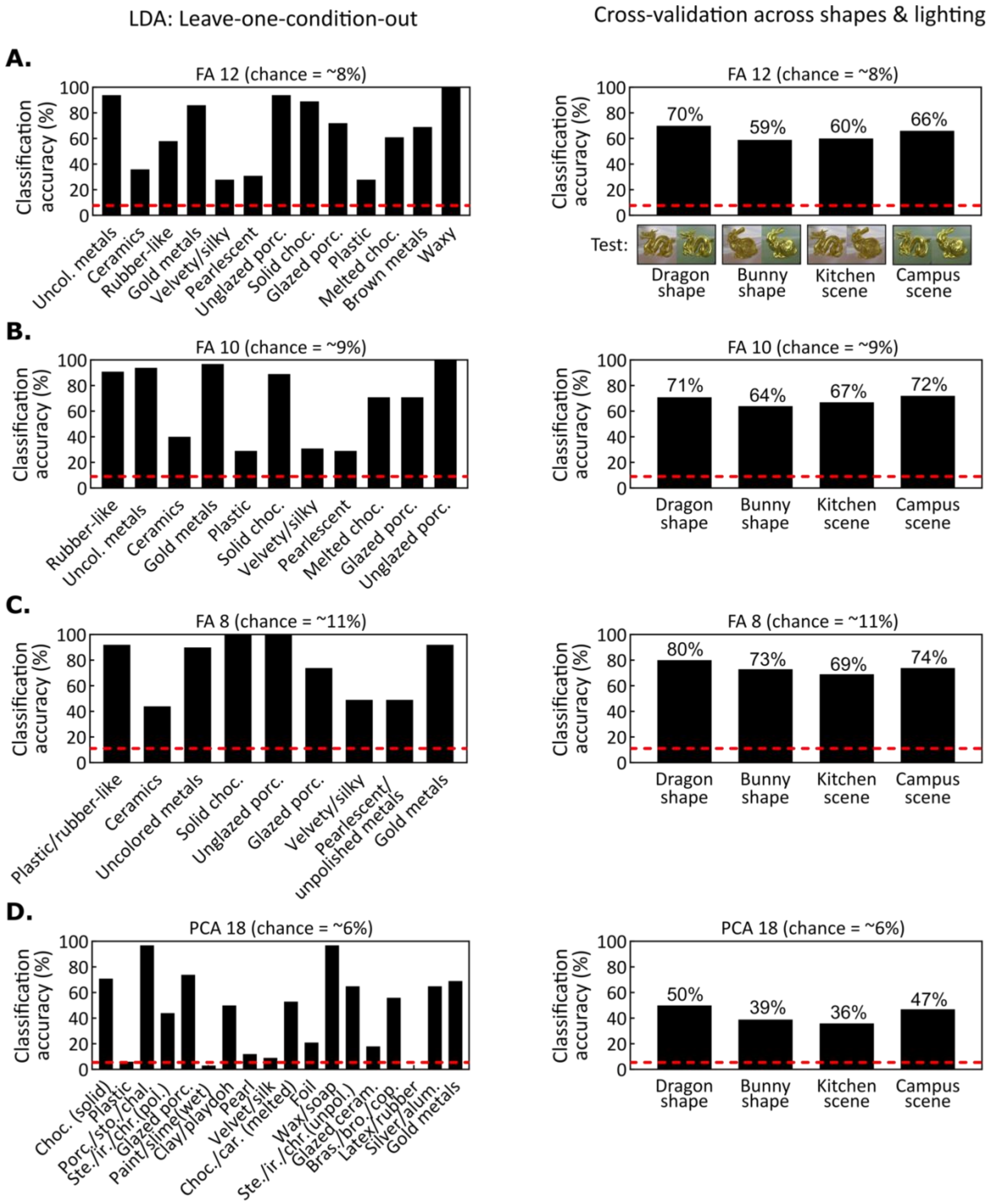


**Supplementary Figure 13.** Results of the linear discriminant analysis with leave one out validation for different factor solutions. Overall we see the similar classification performance as in the 12 factor solution shown in Figure 6B and panel A.

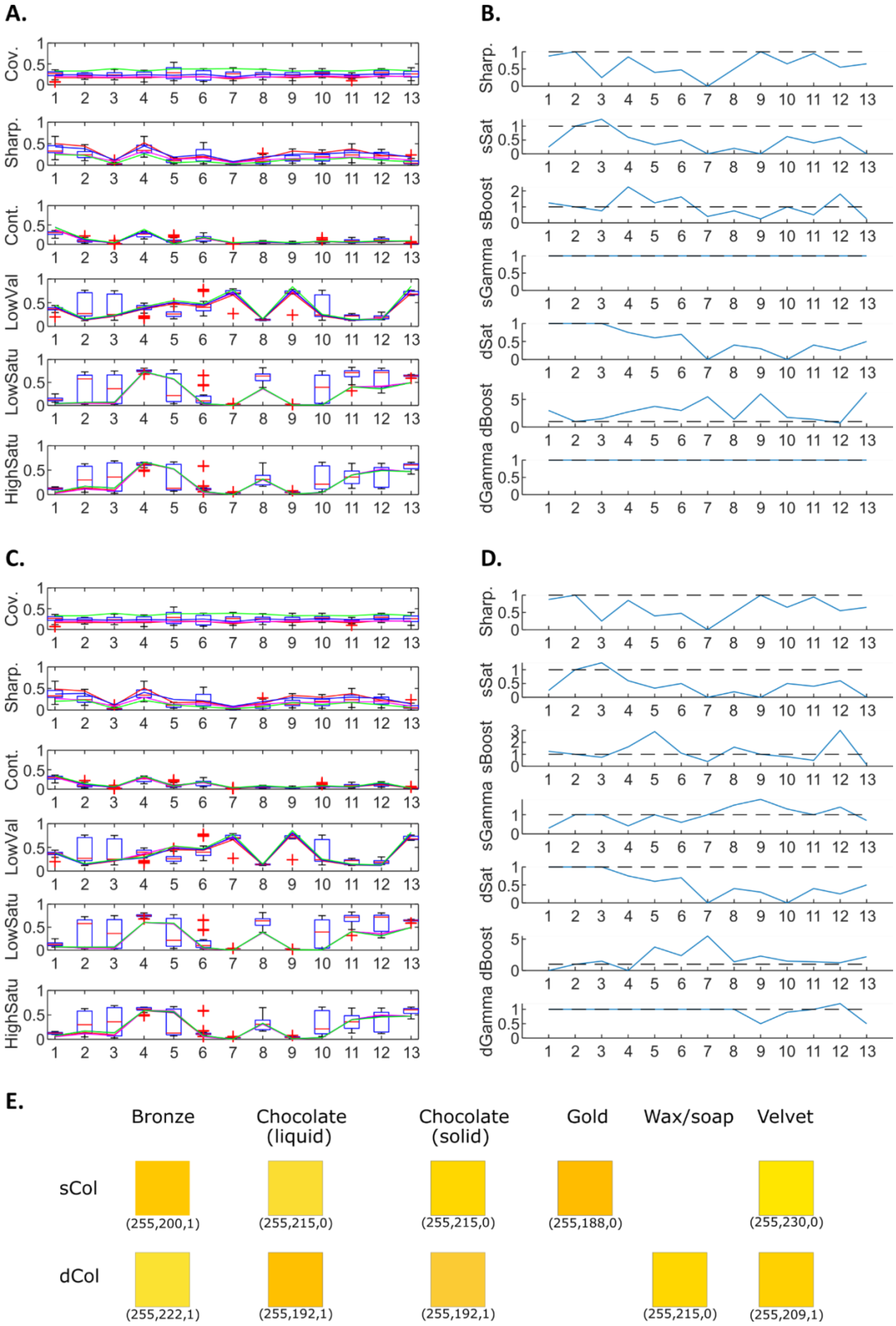
A.
B.
C.
D.
E.
Cov.
Sharp.
Cont.
LowVal
LowSatu
HighSatu
sSat
sBoost
sGamma
dSat
dBoost
dGamma
Bronze
Chocolate (liquid)
Chocolate (solid)
Gold
Wax/soap
Velvet
sCol
dCol
(255,200,1)
(255,215,0)
(255,215,0)
(255,188,0)
(255,230,0)
(255,222,1)
(255,192,1)
(255,192,1)
(255,215,0)
(255,209,1)

**Supplementary Figure 14.** Information about how the visual gloss features were manipulated in Experiment 4. Numbers on x-axis are the emergent categories (ordered as in Figure 4A in the manuscript). Note that 2 is the original glazed ceramic stimuli that the manipulations were performed on. (A) and (C) show measurements of visual gloss features for the simple feature manipulations and complex feature manipulations, respectively. The box plots visualize the distribution of measured gloss features for rendered stimuli (i.e., they show the same data than the violin plots of Figure 6A in the manuscript, organized per feature instead of category), with '+' symbols representing outliers. The lines visualize the gloss features measured for the manipulated stimuli, with one line for each of the 4 scenes (red for Dragon/Kitchen, magenta for Bunny/Kitchen, blue for Dragon/Campus, green for Bunny/Campus). Note how these latter measured features pass through the box plots for each category, with very similar feature values across different scenes. The features varying the most across scenes are sharpness and coverage, which are logically more affected by different lighting and geometry. (B) and (D) show the image manipulation parameters for the simple feature manipulations and complex feature manipulations, respectively. The dashed horizontal lines correspond to the default values for each parameter; note that for category number 2 (i.e., glazed ceramic), all parameters have the default value since we start from this category. Moreover, the gamma parameters for the specular and diffuse components remain at the default value of 1 for the simple feature manipulations in (B). In (E), we show the materials for which the diffuse and/or specular component colour have been pre-multiplied. We systematically chose bright and saturated colours for this step and relied on intensity and saturation adjustments to obtain more desaturated colours, as can be seen in (B) and (D).

**A.**

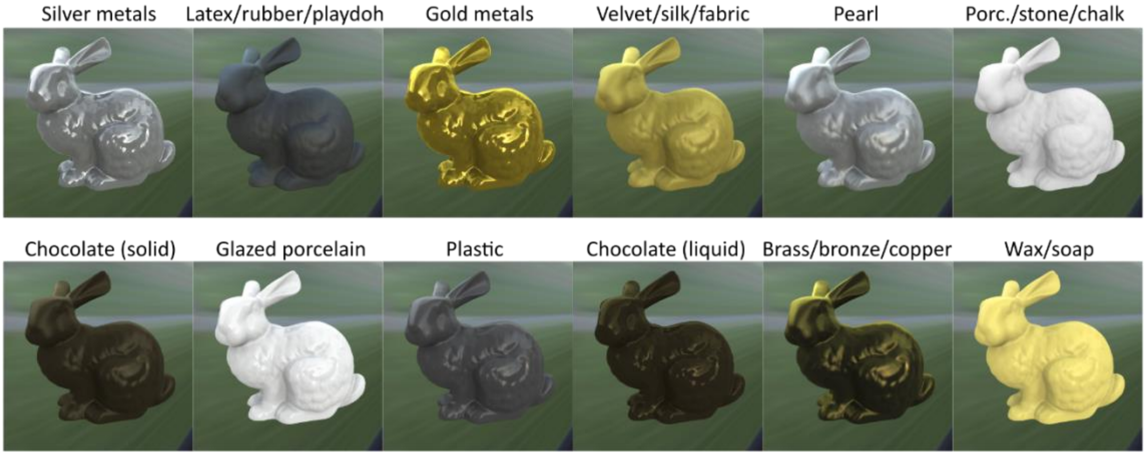


**B.**

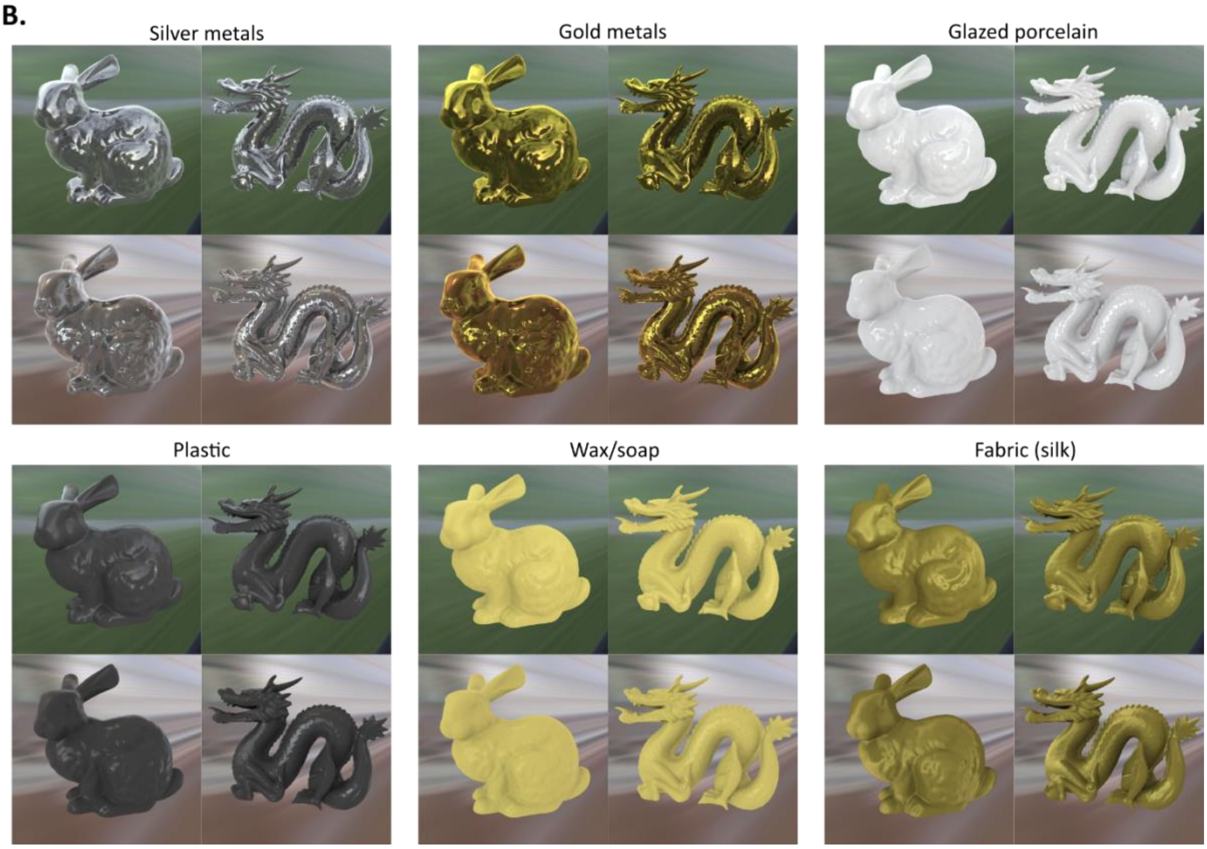


**Supplementary Figure 15.** Example stimuli from the feature manipulation experiment (Experiment 4), labelled with the intended categories. **A.** Examples from the simple feature manipulations on a single scene (bunny object, campus lighting). **B.** Examples of the effects of adding the complex feature manipulations for all four scenes. Using gamma remapping permits to raise the intensity of lowlight reflections in silver and gold manipulations. A similar gamma remapping permits to soften the shadows of the diffuse component in the glazed porcelain, plastic, and wax manipulations, hence mimicking diffusion inside the material. The specific filter for silk fabric creates pairs of elongated highlights with sharp boundaries, which mimics the effect of anisotropy found in such materials.

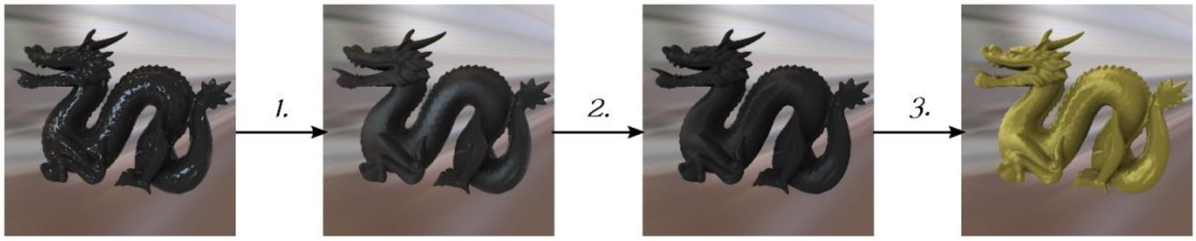


**Supplementary Figure 16.** Image manipulation process used to mimic a satin/velvet appearance (Experiment 4). 1. The specular component of the input image (“Glazed ceramic”) is blurred using a large kernel. 2. A non-linear intensity remapping is applied to the specular term only; it has the effect of darkening the centre of highlights, creating elongated highlights with sharper borders. 3. The colour and intensity of the diffuse and specular components are adjusted as with other material manipulations.

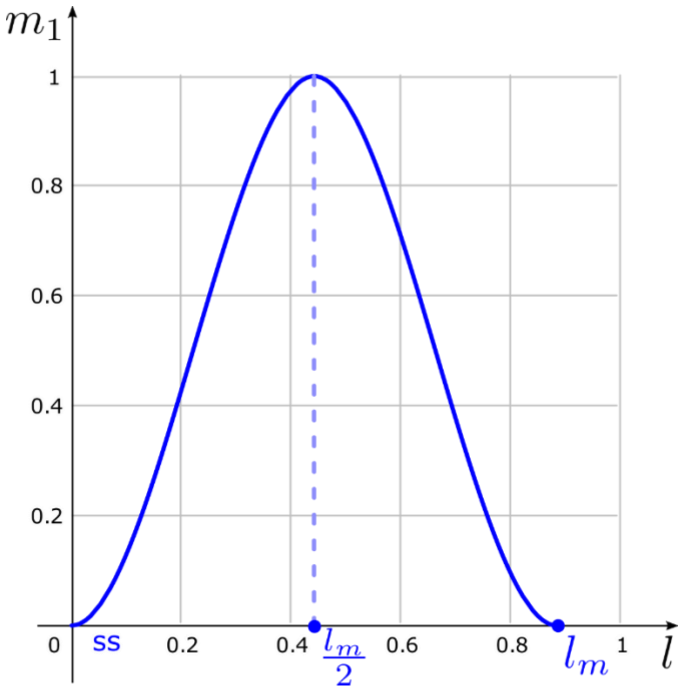


**Supplementary Figure 17.** The non-linear intensity remapping that was applied to the specular term during feature manipulations (Experiment 4).
